# Condensation tendency of connected contractile tissue with planar isotropic actin network

**DOI:** 10.1101/2020.06.23.168237

**Authors:** Tianfa Xie, Sarah R. St. Pierre, Nonthakorn Olaranont, Lauren E. Brown, Min Wu, Yubing Sun

## Abstract

It has been found that many types of cells form nematic symmetry on confined planar substrates. Such observation has been satisfactorily explained by modeling cells as crowded self-propelled rods. In this work, we report that rat embryonic fibroblast (REF) cells when confined in circular mesoscale patterns, form a new type of symmetry where cells align radially at the boundary. Unlike NIH-3T3 and MDCK monolayers, the REF monolayer presents a supracellular actin gradient with isotropic meshwork. In addition, the contractile REF cells present strong adhesive interactions with neighboring cells, which confers the monolayer with significant condensation tendency. We found the loss of condensation tendency by inhibiting the cell contractility or disrupting cell-cell adhesion led to the disappearance of the radial alignment. In theory, we found the prestretch due to condensation tendency with differential cell stiffness is sufficient to explain the new symmetry within a confined tissue continuum.

## INTRODUCTION

The collective migration and rearrangement of cells plays a critical role in various biological processes such as morphogenesis^1^, wound healing^2^, and cancer metastasis^3^. Mechanical variables including cell-cell adhesion forces, cell-substrate traction forces, and self-propulsion have been used to describe the mechanical behavior of a cell monolayer^4^. Using this framework, the collective behaviors of epithelial-like cells with well-defined cell-cell junctions and cortically presented cytoskeleton have been successfully described^5–7^. Recently, theories of active nematic liquid crystals have been applied to monolayers of spindle-like cells such as NIH-3T3 fibroblasts^8^ and neural progenitor cells^9^. In these works, individual cells are treated as incompressible, elongated, and self-propelled particles, and the long-range alignment of these cells is well described by the equilibrium liquid crystal theory. The same approach has also been expanded to epithelial cells with polarized actin cytoskeleton, and reveals that cells located at topological defects experience large compressive stress which leads to the apoptotic exclusion of cells^10^.

A general observation obtained in these nematic-like cell systems is that cells align their shapes with one another, and eventually form topological defects with half-integral winding numbers^8–10^. When confined in patterned mesoscale circular islands, the cells on the boundary of patterns always align circumferentially^8^. This has been demonstrated as a general characteristic of many cell types such as retinal pigment epithelial cells, mouse myoblasts, melanocytes, osteoblasts, and adipocytes^8, 11^. Active nematic theory has been also applied in multiscale systems ranging from polar filament to school of fish^12^. Notably, in addition to the nematic phase, polarized motor proteins display spiral or aster patterns *in vitro*, resembling the polar phase of liquid crystal^13^. In multi-cellular systems, it has been found that skeletal muscle cells with anisotropic actin bundles can align along the direction of actin stress fibers to form a cell sheet *in vitro* guided by geometrical cues^14^.

Distinct from skeletal muscle cells or spindle-like fibroblasts, various mesenchymal-like cells such as neural crest cells^15^, mesenchymal stem cells^16^, and chondrocytes^17^, express a significant level of intercellular adhesion molecules like N-cadherin, and have isotropic actin network. While the collective cell behaviors are critical for their biological functions such as collective migration and condensation, it is unclear how the mechanical interaction among these cells contribute to their collective behaviors and biological functions.

Here we report a unique cell condensation tendency (*i.e.*, reduce in cell area on planar substrates), in cells with strong cell contractility, adhesion, and isotropic actin network. Using a rat embryonic fibroblast cell line (REF-52), which, unlike 3T3 fibroblasts, express a significant level of cell adhesion molecules such as N-cadherin and β-catenin^18^, we show that these cells can robustly self-organize into polarized organization when confined in circular mesoscale patterns on a rigid substrate. In contrast to incompressible 3T3 fibroblasts forming the nematic phase, REF cells on the boundary of circular patterns show significant area expansion and radial alignment. Using a continuum model and a Voronoi cell model, we discover that such condensation tendency, together with an autonomously established supracellular actin gradient, are sufficient to explain the observed radial alignment in confined circular geometries. The distinct behavior of REF-like cells may play a functional role in the collective migration and condensation, which are conventionally considered driven by chemotaxis^19, 20^.

## MATERIALS AND METHODS

### Cell culture

Original Rat Embryo Fibroblast cell line (REF-52) stably expressing yellow fluorescent protein (YFP) - paxillin fusion protein is a gift from Dr. Jianping Fu. REF 11b and 2c subclones were generated by single-cell clone selection. Cells were maintained in high-glucose Dulbecco’s modified Eagle’s medium (DMEM; Invitrogen) supplemented with 10% fetal bovine serum (FBS; Invitrogen), 4 mM L-glutamine (Invitrogen), 100 units/mL penicillin (Invitrogen), and 100 μg/mL streptomycin (Invitrogen). NIH/3T3 cells (a gift from Dr. Mingxu You) were cultured in DMEM medium supplemented with 10% calf serum, 100 units/mL penicillin (Invitrogen), and 100 μg/mL streptomycin (Invitrogen). Both 3T3 and REF cells were subcultured at about 80% confluency following standard cell culture procedures. All cells were cultured at 37 °C and 5% CO_2_.

### RNA sequencing and data analysis

Total RNA was extracted from REF subclones 2c and 11b using the Aurum Total RNA Mini Kit (Bio-rad) following the manufacturer’s instructions. RNA quality was assessed using 6000 Nano Agilent 2100 Bioanalyzer (Agilent Technologies, Santa Clara, CA). The concentration of the libraries was measured using Qubit 3.0 fluorometer (Life Technologies, Carlsbad, CA). cDNA libraries were single-end sequenced in 76 cycles using a NextSeq 500 Kit v2 (FC-404-2005, Illumina, San-Diego, CA). High-throughput sequencing was performed using NextSeq500 sequencing system (Illumina, San-Diego, CA) in the Genomic Resource Laboratory of the University of Massachusetts, Amherst. All sequencing data were uploaded to the GEO public repository (https://www.ncbi.nlm.nih.gov/geo/) and were assigned series GSE148155. Validation of sequence quality was performed using the BaseSpace cloud computing service supported by Illumina (BaseSpace Sequence Hub, https://basespace.illumina.com/home/index). RNA-seq reads were aligned to the rat reference genome (Rattus norvegicus UCSC rn5) using TopHat Alignment. Then, the differential gene expression analyses were performed by Cufflinks Assembly & DE using previous alignment results produced by the TopHat app as input. Shortlists of significantly differentially expressed genes were identified by applying thresholds of 2-fold differential expression and false discovery rate q ≤ 0.05.

### Cell migration assay

To track the migration of patterned REF 2c cells, brightfield live-cell imaging was performed at 37°C, 5% CO_2_ using an automated digital microscope with a 10× objective with a gas controller (Cytation 3 microplate reader, BioTek Instruments Inc., Winooski, VT, USA). Images were collected every 10 minutes for 24h. Acquired brightfield images were merged and corrected for frame drift. To analyze cell migration, individual cell positions were manually tracked using CellTracker software^21^ implemented in MATLAB (MATLAB R2020a, MathWorks). The average speed for each cell was calculated as the total migration length of each cell divided by the total time. Mann–Whitney test was used to compare the migration length and average speed of the cells since the data were not normally distributed (Shapiro-Wilk test). Statistical differences were defined as where *, *P* < 0.05; **, *P* <0.01; ***, *P* <0.001.

### Microcontact printing

Soft lithography was used to generate patterned polydimethylsiloxane (PDMS) stamps from negative SU8 molds that were fabricated using photolithography. These PDMS stamps were used to generate patterned cell colonies using microcontact printing, as described previously^22^. Briefly, to generate patterned cell colonies on flat PDMS surfaces, round glass coverslips (diameter = 25 mm, Fisher Scientific) were spin-coated (Spin Coater; Laurell Technologies) with a thin layer of PDMS prepolymer comprising of PDMS base monomer and curing agent (10:1 *w*/*w*; Sylgard 184, Dow-Corning). PDMS coating layer was then thermally cured at 110 °C for at least 24 h. In parallel, PDMS stamps were incubated with a fibronectin solution (50 μg·ml^−1^, in deionized water) for 1 h at room temperature before being blown dry with a stream of nitrogen. Excess fibronectin was then washed away by distilled water and the stamps were dried under nitrogen. Fibronectin-coated PDMS stamps were then placed on top of ultraviolet ozone-treated PDMS (7 min, UV-ozone cleaner; Jetlight) on coverslips with a conformal contact. The stamps were pressed gently to facilitate the transfer of fibronectin to PDMS-coated coverslips. After removing stamps, coverslips were disinfected by submerging in 70% ethanol. Protein adsorption to PDMS surfaces without printed fibronectin was prevented by incubating coverslips in 0.2% Pluronic F127 solution (P2443-250G, Sigma) for 30 min at room temperature. Coverslips were rinsed with PBS before placed into tissue culture plates for cell seeding. For patterned cell colonies, PDMS stamps containing circular patterns with a diameter of 344 μm, and ring patterns with outer diameter 400 μm and inner diameter of either 200 μm (thick ring) or 300 μm (thin ring) were used.

### Immunocytochemistry

4% paraformaldehyde (Electron Microscopy Sciences) was used for cell fixation before permeabilization with 0.1% Triton X-100 (Fisher Scientific). Cells were blocked in 10% donkey serum for 1 h at room temperature. Primary antibodies used were beta-catenin from rabbit and anti-α-tubulin from mouse (Proteintech). For immunolabelling, donkey-anti goat Alexa Fluor 488, donkey-anti rabbit Alexa Fluor 555, donkey-anti mouse Alexa Fluor 647. For actin microfilaments visualization, Alexa Fluor 488 conjugated phalloidin (Invitrogen) was used. Samples were counterstained with 4,6-diamidino-2-phenylindole (DAPI; Invitrogen) to visualize the cell nucleus.

### Traction force measurement

The protocol for generating microposts and measuring traction force has been published previously^23^. For single cell traction force measurement, REF cells were incubated for 48h on DiI stained micropost arrays, then live-cell imaged. Microposts with a diameter of 2 μm and a height of 8.4 μm (effective modulus *E_eff =_* 5 kPa) were used. Custom MATLAB script was written to quantify post deflection using Eq. 1:

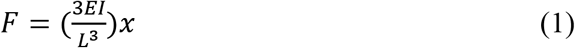

where *F* is the force applied to the tip of the post, *E* is the elastic modulus of PDMS, *I* is the area moment of inertia, *L* is the post height, and *x* is the deflection of the post tip (MathWorks; https://www.mathworks.com/).

### Image analysis

Phase contrast and fluorescence images of patterned cell colonies were recorded using an inverted epifluorescence microscope (Leica DMi8; Leica Microsystems) equipped with a monochrome charge-coupled device (CCD) camera. Since the cell colonies were circle-like in shape and the approximate radii of the circles were known, the centers of the colonies could be found using the circle Hough transformation which is the MATLAB function *imfindcircles*. The distance vector between the center of each cell colony and the center of the image frame shifted each pixel of the image. Using the shifted images, the stacked images could be generated by adding the values at same position of the pixels. The fluorescence intensity of each pixel in stacked images was normalized by the maximum intensity identified in each image. To plot average intensity as a function of distance from the pattern centroid, the stacked intensity maps were divided into 310, 186, and 91 concentric zones for the circle, thick ring, and thin ring, respectively, with single pixel width. The average pixel intensity in each concentric zone was calculated and plotted against the normalized distance of the concentric zone from the pattern centroid.

### Fiber angle deviation

The vector module of the Orientation J plug-in^24^ was used with Fiji^25^ to quantify the angle deviation of actin and α-tubulin stained fibers in REF cell images. Five images were analyzed for each group (circle, thick ring, thin ring). The mean fiber angle deviation was plotted versus the normalized distance from the center of the pattern. A step size of 172/31 ≈ 5.55 μm for the circle, 100/18 ≈ 5.56 μm for the thick ring, and 50/9 ≈ 5.56 μm for the thin ring patterns was used to generate approximately the same number of points respective to the size of the pattern. The schematic of (α), fiber angle deviation from the radius, was drawn in **Figure 4 – figure supplement 1**.

### Structure tensor

The structure parameter, *k_H_*, was calculated from histograms developed from the variation of fiber angle deviation at each concentric distance from the center or innermost edge of the patterns (**Figure 4 – figure supplement 1**). We used a 2D structure tensor *H* to quantify the averaged fiber orientation of the fibers at each point along the radius^26^:

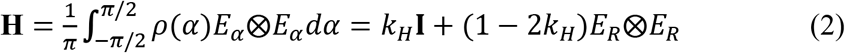

where *α* is the angle between the outward radial direction *E*_*R*_ and actin fiber direction *E*_*α*_ (See **Figure 4 – figure supplement 1**) and *ρ*(*α*) is the normalized orientation density function satisfying *ρ*(*α*) = *ρ*(−*α*) and 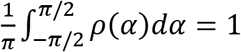. As *I* is the 2 × 2 identity matrix and 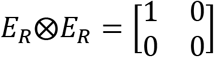, the structure tensor H can be represented by the structure parameter

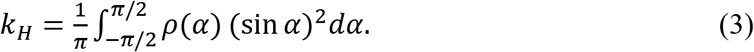

By definition, 0 ≤ *k_H_* ≤ 1, and the fiber distribution is more aligned with the radial (angular) direction as *k_H_* decreases (increases). *k_H_* = 0.5 indicates that the actin fiber distribution is not aligned with either direction. From the actin intensity field, our quantification shows that *k*_*H*_ is approximately 0.4 ~ 0.6 along the radius, suggesting an isotropic distribution of the fiber orientation (**Figure 4 – figure supplement 1D**).

### Continuum mathematical model

#### Tissue elasticity

We assume the elastic stresses dominate over other stresses. We define the energy (per stress-free configuration volume) *W (λ_1_, λ_2_, λ_3_)* as a function of the principle stretches *λ_i_*, *i* = 1, 2, 3, in full 3D. Treating the 2D radially symmetrical multicellular tissue as elastic media reinforced by actin fibers^26^, we have

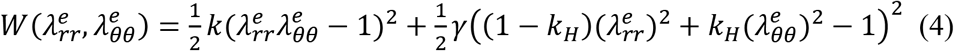

where 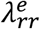 and 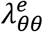 are the principle elastic stretch along the radial and angular direction, respectively, *k* is the bulk elasticity, and *γ(r)* is the local stiffness reinforced by fibers, which is assumed to be positively correlated with the local actin intensity. From experimental data, it is approximately shown that the orientation of the actin fibers is isotropic almost everywhere. As such, we assume *k_H_* = 0.5 along the pattern radius (see **Figure 4-figure supplement 1**) and the hyperelastic energy of the REF pattern to be

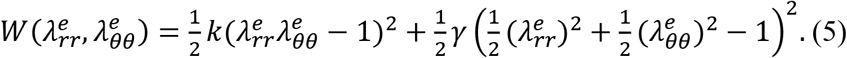

Then we can derive the Cauchy stresses^27^ as 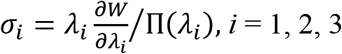,

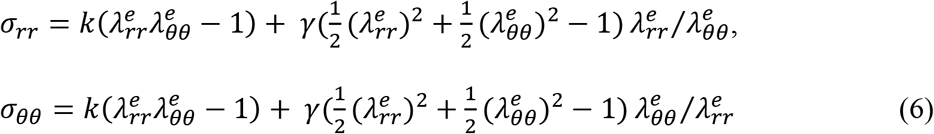

#### Tissue prestretch

We assume that the prestretch due to the condensation tendency is isotropic in all directions, and define 0 < *g* ≤ 1 as the prestretch. *g* decreases as the level of condensation tendency increases. When *g* = 1, there is no condensation tendency. Following the theory of morphoelasticity^28^ in Eulerian frame^28, 29^, we define the elastic stretches 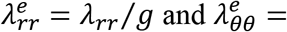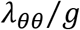 as a result of the mismatch between the prestretch *g* and the observable deformation stretches 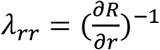 and 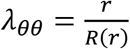. Notice we define a map *R(r)* from the current position to the initial position to track the deformation inversely. This is different from the traditional morphoelasticity which defines a nonlinear deformation map *r*(*R*) from the initial position to the current position. The traditional framework does not allow straightforward coupling between spatially-dependent variables in the lab coordinates, such as the local stiffness *γ(r)*, and the deformation map *r*(*R*) and the associated stresses, which are all defined in the Lagrangian coordinates. Recently we have reframed the morphoelasticity theory by introducing the reference mapping techniques in Ref. 29. In the new framework^29^, all field variables are in the lab coordinates, including the stresses as defined below:

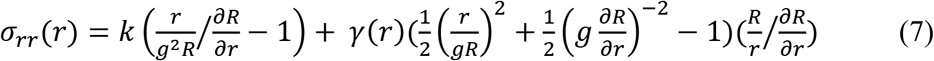

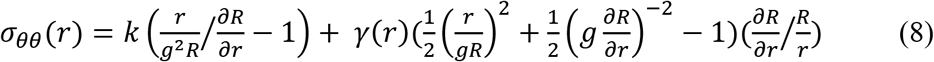

#### Mechanical equilibrium in the microtissue

We assume the elastic stresses dominate over other forces, such as the mechanical interaction with the substrate. Using radial symmetry, we have the force balance equation for the radial and circumferential stress, *σ_rr_* and *σ_θθ_*, respectively:

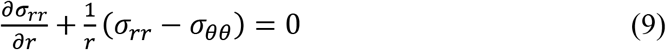

#### Geometric constraint at the boundary

Consider a ring pattern with inner radius *r_in_* and outer radius *r_out_* = 1, we have

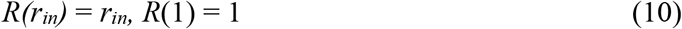

from the experimental observation. All the results (see Figure 4 and the text) are shown by solving Eq. (7), (8) and (9), with the boundary condition (10), and assuming the bulk modulus *k*~0 is negligible compared to the fiber-reinforcing stiffness *γ(r)*.

### Voronoi cell mathematical model

#### Cell-tissue configurations

We begin with a 2D domain and generate polygonal cells from the 2D Voronoi tessellation to represent the cell configurations. The graph of Voronoi cell tessellation is dual to the Delaunay triangulations. In particular, the relation between a trio of adjacent Voronoi cell centers (<**r**_i_, **r**_j_, **r**_k_>) and its corresponding vertex (**ω**_<i,j,k>_) is given by the following equations^30,31^:

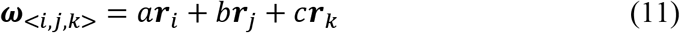

where

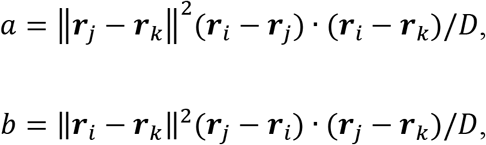

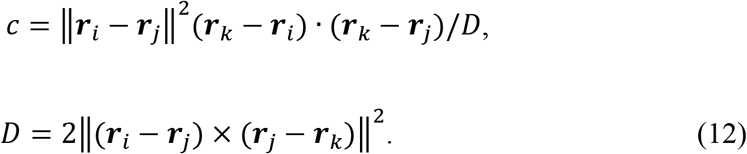

#### Cell prestretch and differential stiffness

To describe the monolayer mechanics, we define a total energy that is generally a function of the areas of cells *A*^*α*^ and lengths of junctions 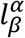:

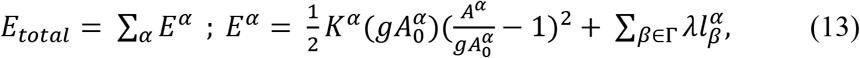

where *α* indicates each cell, *β* indexes junctions of cell *α*, and Γ is the set of junctions of cell *α* with tension. For modeling epithelial cells, the tension is considered on each intercellular junction to represent the net mechanical effect of actomyosin contraction and intercellular adhesion at the apical surface of the tissue. We apply tension to the intercellular junctions, and on the junctions located at the boundaries of the micropattern to ensure its circularity (see *Geometric constraints* below). Similar to the continuum model, we introduce the prestretch 0 < *g* ≤ 1 to describe the condensation tendency to shrink the intrinsic cell size 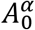 without any mechanical interactions. *g* is applied to all of the cells in the pattern. To describe stiffness gradient between the boundary cells and interior cells, similarly to the continuum model, we introduce the stiffness differential parameter 0 < *ρ* ≤ 1, which is the ratio of the stiffness *K*^*α*^ between boundary and interior cells. *ρ* is only applied for the cells on the boundary of the pattern.

#### Mechanical equilibrium among cells

*E*_*total*_ is a function of the coordinates of vertices via its dependence on lengths of junctions and areas of cells. In particular, the length of each junction is 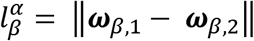 where 1 and 2 indicate the adjacent vertices of *β* junction, and the area of each cell is 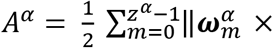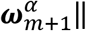, where *Z*^*α*^ is the number of vertices of cell *α* (Notice that 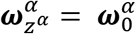. In addition, the coordinates of vertices depend on the coordinates of a trio of neighboring Voronoi cell centers through Eqs. (11) and (12). By chain rule, we can solve the cell configurations by minimizing the energy *E*_*total*_ following the dynamics of cell centers:

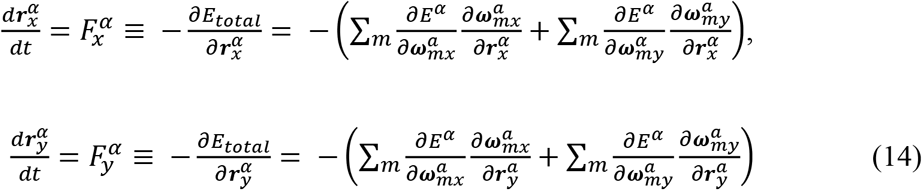

where *m* is the index of the vertices of cell *α*. The subscripts x and y indicate the component x or y of the vector we are considering.

#### Geometric constraint at the boundary

We define a full circle or annulus inside the squared domain to initialize the micropattern, by calibrating the size ratio between individual cell and the micropattern according to the *in vitro* setup. Initially, there are 625 cells in the squared domain. Then, the circular or annular domain defined by thresholding the distance of cells from origin does not provide the circular borders. To round up the borders, we minimize the following energy

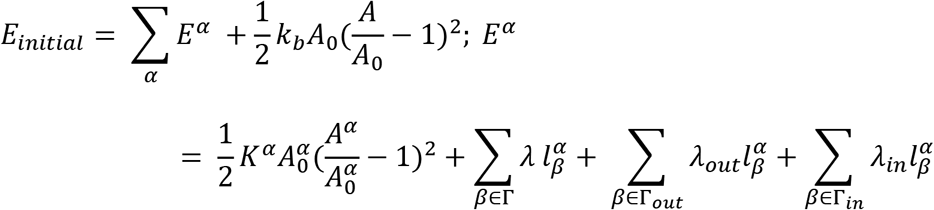

with *K*^*α*^ = 1, *λ* = 15, *λ*_*out*_ = 20, *λ*_*in*_ = 5, and *k*_*b*_ = 0.1. Then tension terms 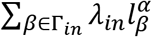 and 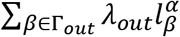 are included to ensure the circularity of the inner and outer boundaries, respectively, and the single term 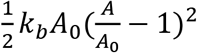 is included to ensure that the total area *A* of the micropattern is close to the total initial area *A*_0_. Once *E*_*initial*_ reached a local minimum, the centroids of the boundary cells and cells outside of the circular or annulus domain were fixed. When the prestretch, stiffness differential, and tension on the intercellular junctions are considered in Eq. (13), all the non-boundary cell centers move and reach equilibrium following Eq. (14).

### Statistical analysis

Student’s t-test was used when there were two groups. One-way ANOVA and post-hoc Tukey’s test were used for three or more groups. Mann–Whitney test was used for data that was found not normally distributed. Data are represented as mean ± s.e.m.

## RESULTS

### REF 2c cells align radially on circular mesoscale patterns

We first sought to investigate whether various types of cells with different intra- and inter-cellular forces and cell shape have similar self-organization behavior under confinement. We found that when REF 2c cells, a subclone of REF52 cell line, were placed on circular micro-contact printed patterns (diameter = 344 μm), the boundary cells radially aligned over a period of 48 h after cell seeding (**Figure 1A**). Video tracking of individual inner and boundary cells showed that the radial alignment of boundary cells was not caused by oriented cell migration towards or away from the center (**Figure 1 – figure supplement 1A**), as most boundary cells stayed on the periphery of the pattern. Moreover, both the innermost and boundary cells migrated longer and faster between 24-36 h than 36 – 48 h (**Figure 1 – figure supplement 1B-C**). This reduced cell migration coincided with the formation of radial alignment of cells on the pattern boundary. In contrast, 3T3 fibroblasts maintained circumferential alignment on the boundary at both 24 and 48 h (**Figure 1A**), consistent with previous reports. To quantify the cell alignment, we traced the cell outlines and measured the cell angle deviation, defined as the angle between the long axis of an ellipse-fitted cell and the line that connects the pattern center and the centroid of the cell; cell elongation, defined as the ratio of the major axis to the minor axis of the ellipse-fitted cells; and projected cell area using an ellipse-fitting based method (**Figure 1 – figure supplement 2**). We found that for REF 2c cells, the boundary cells were significantly more aligned with the radial direction than the inner cells at 48 h (**Figure 1B**). They were also significantly more elongated and had a significantly larger cell area (**Figure 1C, D**). The boundary 3T3 cells were significantly more circumferentially aligned and elongated than the inner cells but did not have any significant differences in cell area (**Figure 1B-D**). Confocal fluorescence images of REF 2c cells stained with a cell membrane permeable dye CellTracker-Green suggested that the average tissue thickness did not change significantly due to the cell radial alignment, while the boundary cells had a wedge-shape with a drastically reduced cell thickness on the pattern boundary side (**Figure 1 figure supplement 3**).

**Figure 1.**
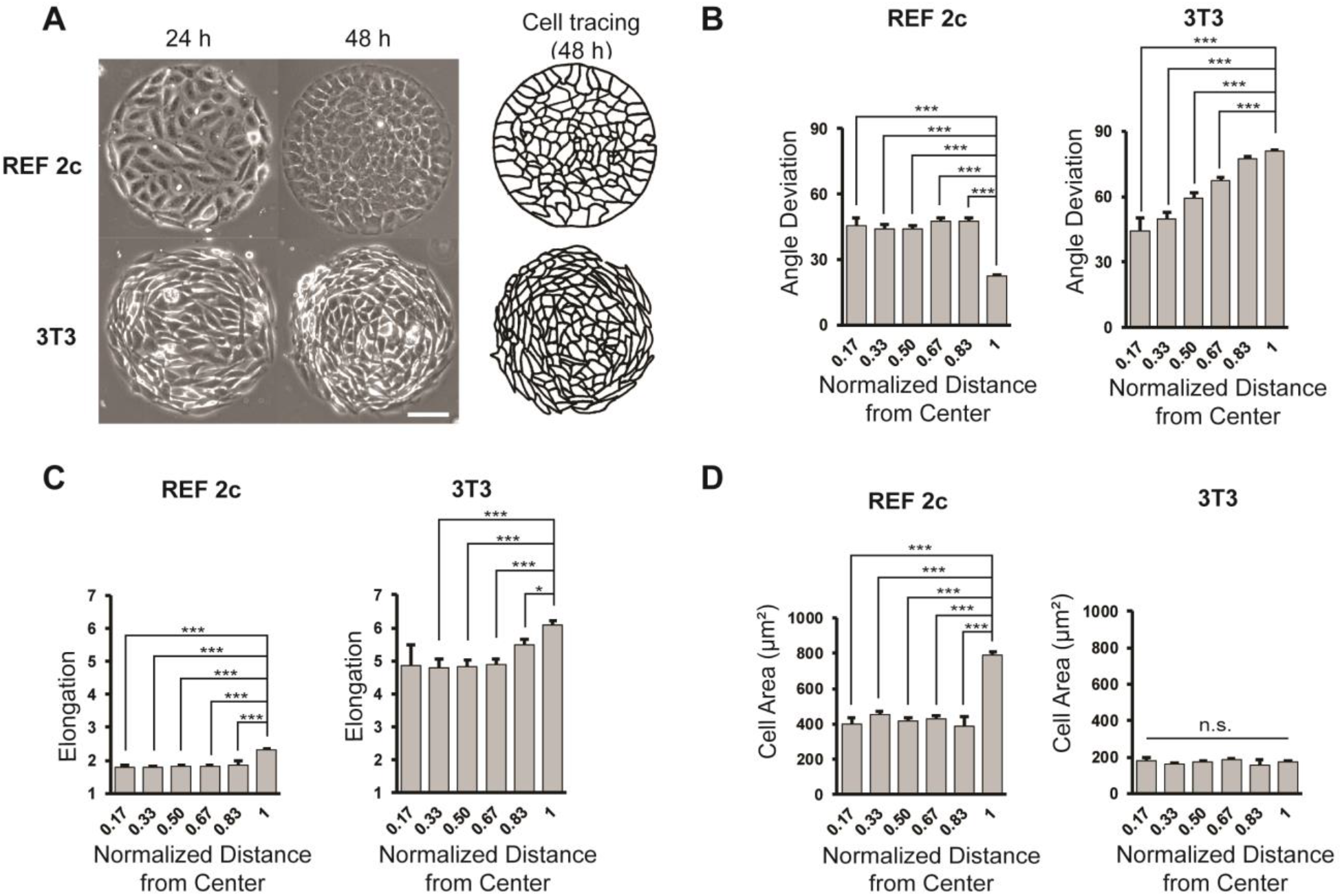
REF 2c cells align radially on circular mesoscale patterns. (**A**) Phase images showing REF 2c and 3T3 cells cultured on patterns for 24 and 48 hrs. Scale bar: 100 μm. Cell tracing showed the cell boundaries at 48 hrs. (**B**) REF 2c and 3T3 cell angle deviation at 48 h as a function of normalized distance from the center of the pattern. *n* = 16 patterns. (**C**) REF 2c and 3T3 cell elongation at 48 h as a function of the normalized distance from the center of the pattern. *n* = 16 patterns. (**D**) REF 2c and 3T3 cell area at 48 h as a function of normalized distance from the center of the pattern. Cell area is defined as the area of the ellipse-fitted cells. *n* = 16 patterns. Data are represented as mean ± s.e.m. *, *P* < 0.05; ***, *P* < 0.001; *n.s.*, *P* > 0.05.

As cells were patterned on PDMS substrates (Young’s modulus *E* = 2.5 MPa), which are significantly stiffer than physiological extracellular matrices, we next tested whether similar phenomenon could be observed on substrates with physiologically relevant substrates. Here we applied a well-established PDMS micropost array (PMA) system with identical surface geometry and different post heights to tune substrate rigidity (**Figure 1 figure supplement 4**)^32^. We found that on soft PMA substrates (*E* = 5 kPa, post height = 8.4 μm), REF 2c cells became polarized at 48 h and condensed towards the center of patterns, resulting in reduced total cell area on each pattern. On stiff PMA substrates (*E* = 1 MPa, post height = 0.7 μm), however, the total cell area on each pattern did not change between 4 to 48 h, and only cells on the boundary became radially aligned, which is consistent with the results on flat PDMS substrates.

### Cell contractility and cell-cell adhesion are required for radial alignment

As all previous works demonstrated circumferential alignment of non-epithelial cells on circular patterns, we asked what factors contribute to the radial alignment of REF 2c cells. We isolated and expanded several subclones of REF cell line, and identified one subclone, named REF 11b, did not radially align at 48 h on circular patterns (**Figure 2A**). RNA-seq data revealed that while most genes have similar expression levels in REF 11b and REF 2c subclones, a small subset of genes were expressed significantly differently (**Figure 2 – figure supplement 1**). As some of these genes are associated with cell adhesion and contraction (e.g., *INTEGRIN* α7 and α8, *MYL9*, and *COL16A1*), we then measured the cell contractility of these two subclones. Traction force measurements for single cells of REF 2c and 11b showed that for both total force and force per area, REF 11b cells were significantly less contractile than REF 2c cells (**Figure 2B**). To verify that the cell contractility was required for radial alignment, we treated patterned REF 2c cells with Blebbistatin and Y27632, drugs known to reduce cell contractility^33^, at 24 h and then imaged the patterned cells at 48 h (**Figure 2C**). The boundary cells of the Blebbistatin and Y27632 treated groups were not significantly more radially aligned than the inner cells, compared to the vehicle control group with DMSO (**Figure 2C**). These results suggest that cell contractility is required for radial alignment formation.

**Figure 2.**
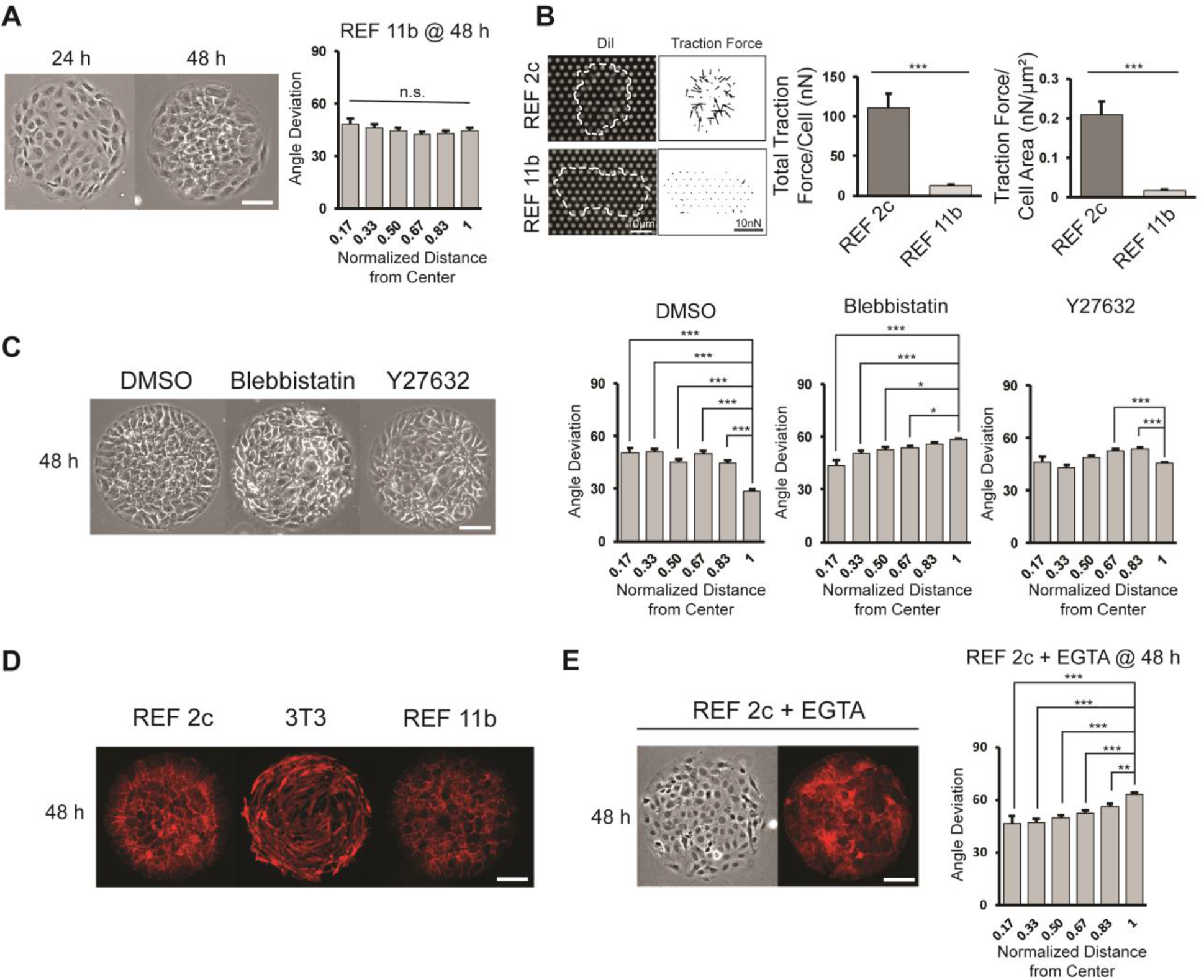
Effect of cell contractility and cell-cell adhesion on cell alignment. (**A**) Phase images of REF 11b at 24 h and 48 h after cell seeding. Average cell angle deviation is quantified with respect to distance from the center of the pattern. *n* = 16 patterns. Scale bar: 100 μm. (**B**) Representative images showing single REF cells cultured on PMA substrates and vector map of deduced traction forces. Plots show the total traction force per cell and traction force per cell area of individual REF 2c and 11b cells. *n* = 15 cells per subclone. (**C**) Phase images of REF 2c treated with DMSO, Blebbistatin, and Y27632 at 48 h. Average cell angle deviation is quantified with respect to distance from the center of the pattern. *n* = 16 patterns per group. Scale bar: 100μm. (**D**) Immunofluorescence images showing the expression of β-catenin in REF 2c, 3T3, and REF 11b cells at 48 h. Scale bar: 100 μm. (**E**) Phase and fluorescence images showing cell orientation and β-catenin expression in EGTA treated REF 2c cells at 48 h. Average cell angle deviation is quantified with respect to distance from the center of the pattern. Scale bar: 100 μm. Data are represented as mean ± s.e.m. *, *P* < 0.05; **, *P* < 0.01; ***, *P* < 0.001; *n.s.*, *P* > 0.05.

As 3T3 cells have a similar level of contractility compared with REF 2c cells^34^, we rationalized that other factors must also contribute to the radial alignment. Staining with β-catenin revealed that REF 2c and REF 11b had clear boundary junctions between cells, while 3T3 cells did not (**Figure 2D**), suggesting such cell-cell adhesion may be required for the establishment of radial alignment. To confirm this, we treated REF 2c cells with EGTA, which reduced cadherin-based cell-cell adhesion without significantly affecting cell adhesion to substrates^35^, at 24 h and then examined the cell alignment at 48 h. We found that when treated with EGTA, the boundary cells were significantly more circumferentially aligned than the inner cells (**Figure 2E**). Together, we identify that cell contractility and cell-cell adhesion are two essential factors required for the establishment of radial alignment of patterned cells. Such tissue level contraction enabled by individual cell contractility and cell-cell adhesion is hereafter termed condensation tendency.

### Supracellular gradient of actin was established in patterned REF cells

Epithelial cells also exhibit significant contractility and strong cell-cell interactions, while previous studies showed that no alignment was found when epithelial cells were confined on similar circular patterns^36^. As a critical difference between epithelial cells and fibroblasts is that actin microfilaments mainly distribute on the cell-cell boundaries in epithelial cell monolayer, while filamentous actin can be found throughout the fibroblast cells. Thus, we next sought to establish the relationship between the spatial distribution of actin microfilaments and the alignment of patterned fibroblasts. Surprisingly, we found a decrease in actin intensity occurred near the boundary of patterns for REF 2c cells, in the same location as the transition between isotropically oriented inner cells and radially aligned boundary cells (**Figure 3**). However, for REF 11b, there was a continued decrease in intensity from the center of the pattern to the outermost edge (**Figure 3**). This is in a sharp contrast to the actin distribution of 3T3 cells, in which no actin gradient was identified (**Figure 3**). Notably, the sharp drop in the intensity profiles near the very end of the intensity plot for actin was an artifact as the outermost cells did not fully cover the pattern boundary.

**Figure 3.**
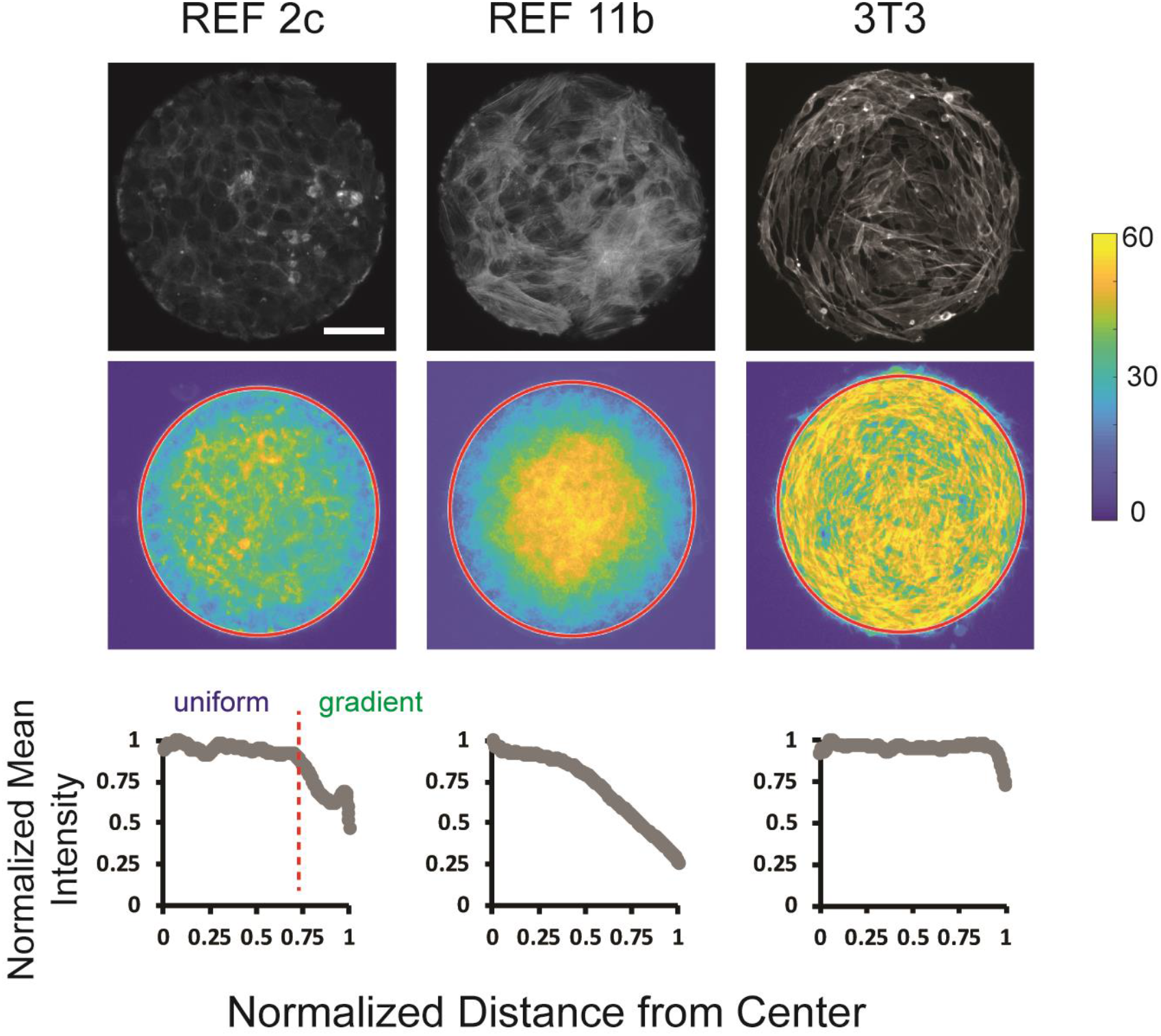
Supracellular actin distribution in patterned fibroblast cells. Top: fluorescence images showing the actin staining of REF 2c, REF 11b, and 3T3 cells. Scale bar: 100 μm. Middle: colorimetric maps showing the actin intensity profiles obtained by overlapping actin staining images. *n* = 20 patterns per group. Bottom: normalized mean intensity of these overlapping images plotted as a function of the normalized distance from the center of the pattern.

### A continuum model demonstrated that the condensation tendency with supracellular actin gradient is sufficient for establishing polarized cell alignment

We next asked what are the deterministic factors that dictate the polar alignment of REF 2c cells. On one hand, the experimental data suggests that the condensation tendency is essential for the polar alignment (**Figure 2**). In addition, the data shows that the alignment is associated with elongation of the boundary cells (**Figure 1**), which suggests that these boundary cells are stretched more along the radial direction than along the circumferential direction. On the other hand, the actin gradient decreasing towards the boundary (**Figure 3**) suggests that there is a gradient of stiffness between the inner and boundary cells, given it is well-documented that actin intensity is proportional to the local cell stiffness^37^. Thus, we hypothesize that the radial alignment and elongation at the boundary is a presentation of the strain field patterned by the condensation tendency and the stiffness gradient. To verify this mechanical mechanism, we derived a continuum mathematical model based on the morphoelasticity theory^26^ (see **Methods**) to study the independent and synergic roles of condensation tendency and stiffness gradient that is challenging to decouple experimentally. Here, we treated the patterned fibroblasts as elastic media reinforced by actin stress fibers, in which elastic stresses dominate over other stresses and mechanical interactions between cell and substrates^38^. Without evidence of the spatially heterogeneity and orientation anisotropy of the cell contractility (see **Figure 1**, **figure supplement 4**), we introduce an isotropic constant prestretch parameter 0 < *g* ≤1 to describe the global condensation tendency in the patterned fibroblasts. The prestretch parameter *g* describes the tendency of the tissue to condensate to g fraction of its area without rigid substrate. *g* =1 corresponds to no condensation tendency. Given the actin orientation being neither radially nor circumferentially aligned (see **Figure 4, figure supplement 1** and **Methods**), we assume that the stiffness of the cells is isotropic, denoted by *γ(r),* and spatially varies as proportional to the local intensity of the actin. For simplicity, we define a linearly decreasing stiffness field 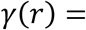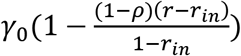 from the micropattern boundary *r* =1 to an interior location 0 ≤ *r*_*in*_ < 1. The stiffness differential parameter, 0 < *ρ* ≤ 1, describes the ratio between the stiffness at the boundary and that in the interior.

**Figure 4.**
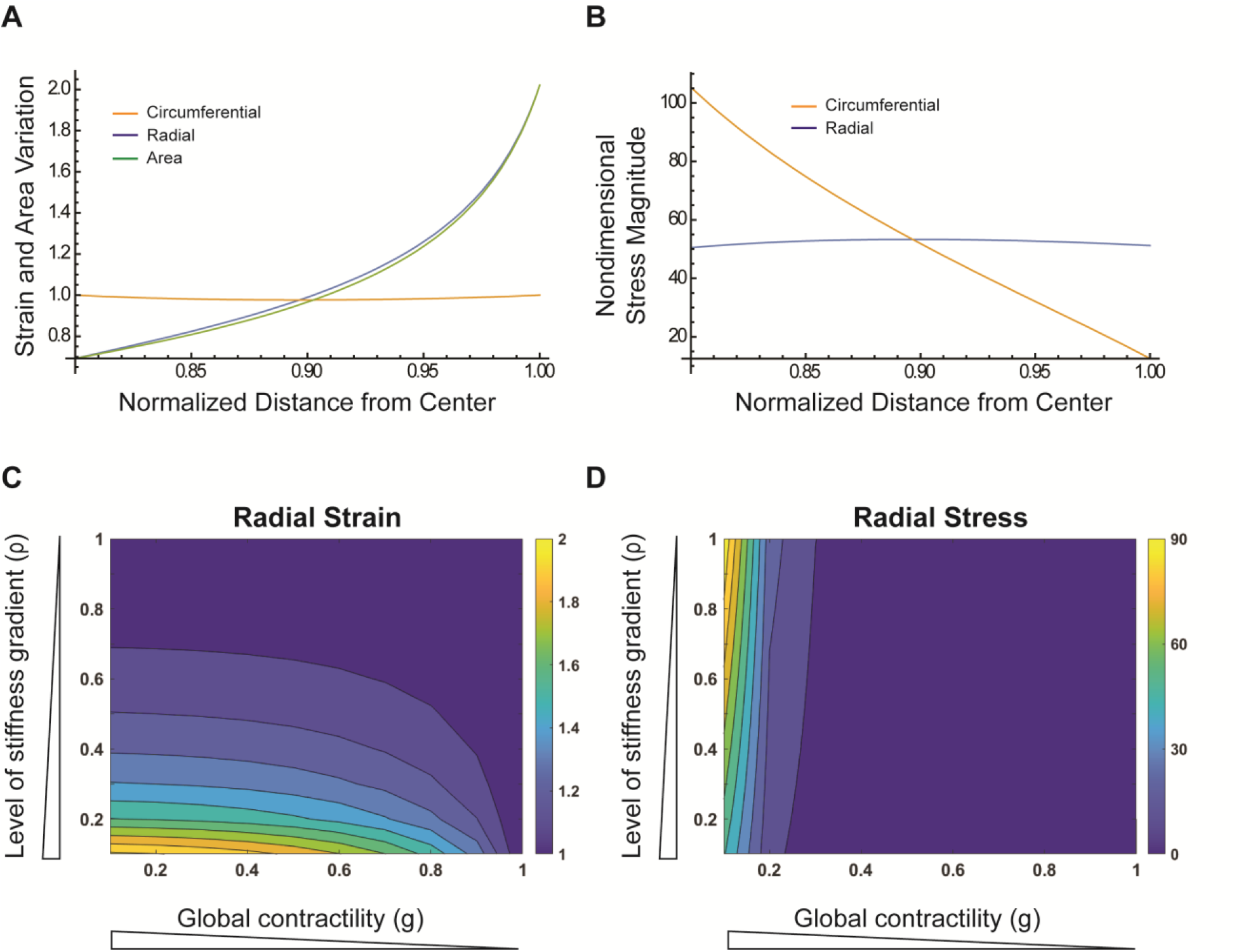
Continuum model of patterned cells. (**A**) Simulation results for strain and area variation as a function of normalized distance from the center (green = area, blue = strain in the radial direction, orange = strain in the circumferential direction). (**B**) Simulation results for stress as a function of normalized distance from the center (blue = stress in the radial direction, orange = stress in the circumferential direction). (**C-D**) Phase diagrams of radial strain (**C**) and radial stress (**D**) calculated from the continuum model as functions of the stiffness differential ρ, and g, the condensation tendency.

Given *g* and *γ(r),* we can solve the deformation stretch field *λ*_*rr*_ and *λ*_*θθ*_ as well as the stress field *σ*_*rr*_ and *σ*_*θθ*_ from Eqs. (7), (8), (9) and (10). When there is no stiffness gradient (*i.e.*, *γ* ≡ *γ*_0_, *ρ* = 0), we have the analytical solution *λ*_*rr*_ = *λ*_*θθ*_ = 1, and *σ*_*rr*_ = *σ*_*θθ*_ = *γ*_0_(1/*g*^2^ − 1). This predicts that without spatial gradient of the stiffness, the condensation tendency will result in a uniform isotropic tensile stress in the radial pattern, and no local deformation associated with cell morphology due to the geometric constraint at the micropattern boundary. Notice when there is no condensation tendency at all (i.e., *g* =1), we have no deformation (*λ*_*rr*_ = *λ*_*θθ*_ = 1) and no stress (*σ*_*rr*_ = *σ*_*θθ*_ = 0) in the tissue continuum, which is the case with the REF 11b cells (**Figure 2B**). When condensation tendency is considered together with stiffness gradient (e.g., *g* = 1/10, *ρ* = 1/10, and *r*_*in*_ = 0.8), the distributions of the strain and stress become nontrivial (**Figure 4A, B)**. From both the condensation experiment (**Figure 1, figure supplement 3**) and the modeling result (**Figure 4A, B**), the cells are all stressed and strained due to the prestretch and are under tensile stresses everywhere along both directions (**Figure 4B)**. While the circumferential tensile stress is lower than the radial stress at the outer boundary, it increases to the level above the radial stress at the interior region. Noticeably, although the stress within the cell colony has been experimentally measured previously^5^, it is challenging in our case because of the extreme tissue condensation on soft substrates. For the strain field, the tissue close to the outer boundary is strained along the radial direction while the interior region is geometrically compressed (**Figure 4A)**. The radial strain directly contributes to an increase of area at the boundary and a decrease of area at the interior. This is consistent with the observation of cell elongation and area at the pattern boundary (**Figure 1C, D**).

We further analyzed the sensitivity of radial stress and radial strain to changes in condensation tendency *g* and the stiffness differential *ρ* at the pattern boundary *r* = 1. Our results indicated that radial strain was more sensitive to changes in the level of stiffness gradient than to changes in the condensation tendency (**Figure 4C, D**). The smaller the *ρ* is (indicating that the exterior stiffness was much small than the interior), the greater the boundary radial strain will be. Radial stress was found to be sensitive to changes in condensation tendency, where a high degree of condensation tendency was required for high level of radial stress to form (**Figure 4D**). Our continuum model revealed that condensation tendency and supracellular stiffness gradient are sufficient for the radial alignment of cells by elongating cells at the boundary, by providing an anisotropic stress field near the tissue boundary which results in a larger radial strain. However, the level of stiffness differential, rather than the level of condensation tendency due to cell contractility and adhesion, is the control parameter for the elongation morphology.

### Voronoi cell modeling predicted REF 2c radial alignment

While our continuum model described the cell elongation in the radial direction governed by cell stiffness gradient and contractility, it did not directly demonstrate that these two mechanical effects are responsible for the observed morphological characteristics of cells including cell area changes, elongation, and angle deviation from the radius in the experiments. Thus, we also considered the condensation tendency with stiffness differential in a Voronoi cell model^30, 31^ to quantitatively compare those characteristics between the theory and the experiments (see **Methods**). Similar to the continuum model, here we implemented a ratio *ρ* between the inner cells and the boundary cells in their stiffness and kept the prestretch *g* uniform. The decrease of both *ρ* and g resulted in the emergence of a distinct boundary cell layer with radial alignment, area increase, and elongation, compared to the inner cells (**Figure 5**). Notice boundary cell radial alignment and area increase, mimicking the *in vitro* phenomenon, do not occur when either contractility or boundary cell stiffness differential was not considered (see the cases with *ρ* = 1 or *g* = 1 in **Figure 5B and 5C, and Figure 5, figure supplement 1A and 1B)**. In summary, we have shown that condensation tendency with stiffness differential near the micropattern boundary is sufficient to replicate the radial alignment and increased elongation and area of the boundary cells compared to inner cells in the REF 2c *in vitro*.

**Figure 5.**
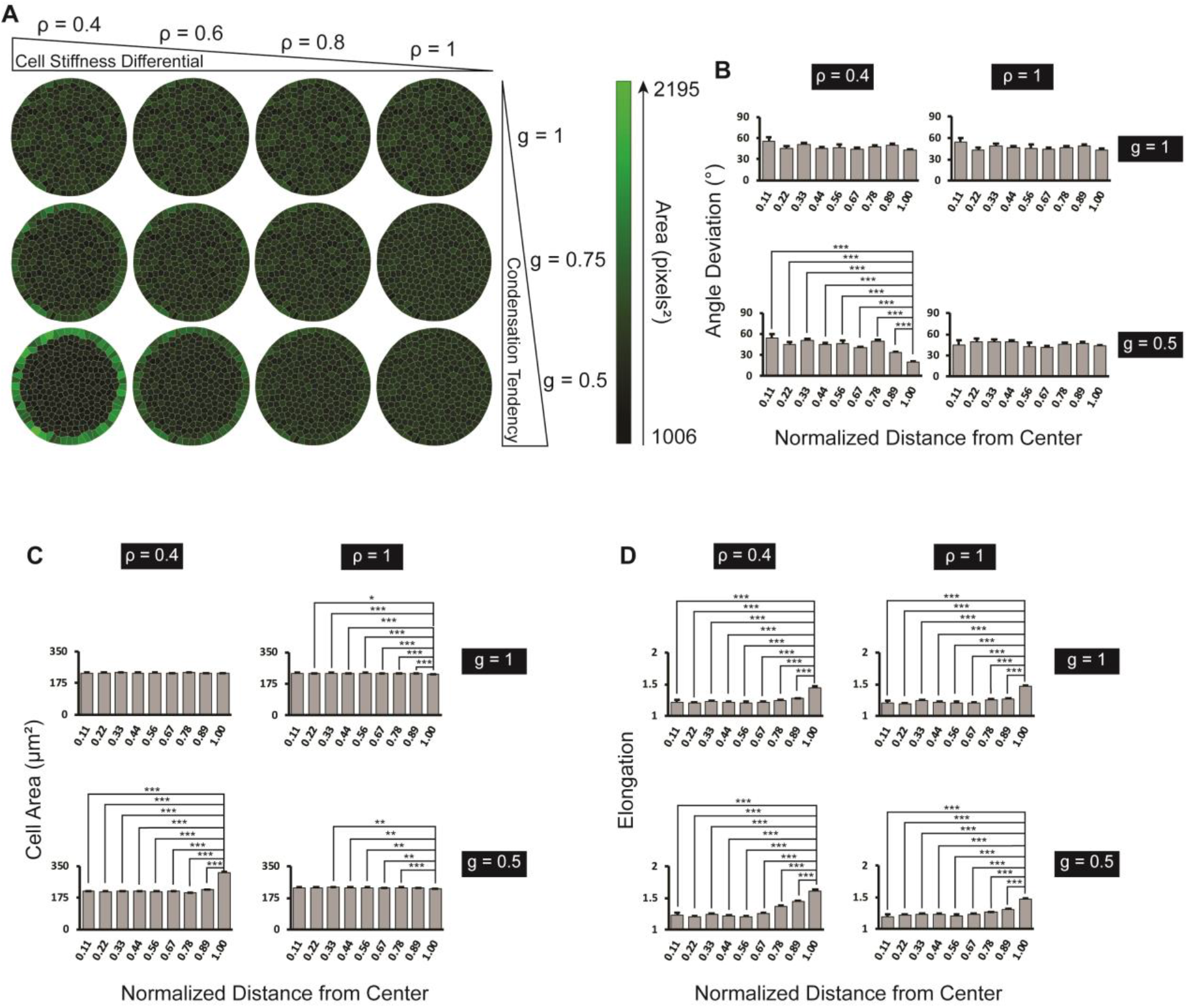
Voronoi cell modeling predicts REF 2c cell behaviors in circular pattern. (**A**) Voronoi cell modeling results from varying condensation tendency and cell stiffness differential at the boundary. (**B**) Angle deviation is plotted as a function of distance from the center of the pattern. *n* = 5 patterns. (**C**) Cell area is plotted as a function of distance from the center of the pattern. *n* = 5 patterns. (**D**) Elongation is plotted as a function of distance from the center of the pattern. *n* = 5 patterns. Data are represented as mean ± s.e.m. *, *P* < 0.05, **, *P* < 0.01, ***, *P* < 0.001.

### Actin gradient and condensation tendency maintain under the change of tissue topology

Our results demonstrated that the emergence of cell stiffness gradient along the radial direction is critical for the cell alignment. We next investigated whether such gradient maintains at the outer boundary under the change of the topology. To do so, we designed two ring patterns with different inner diameters (200 μm and 300 μm) and the same outer diameter (400 μm). We found that surprisingly, both REF 2c and 11b cultured on ring patterns robustly showed an actin intensity gradient from the center to the boundary, regardless of the change of topology and the radius of the inner boundary. In contrast, 3T3 cells did not have an actin gradient for any ring patterns (**Figure 6A, B**). Similar to the circular patterns, we calculated the actin fiber angle deviation and the structure parameter *k_H_* for REF 2c cells (**Figure 6 – figure supplement 1**). Compared to the full circle pattern, the distribution of the structure parameter *k_H_* in ring patterns reveals that the actin fibers are mostly aligned along the tangential direction at the inner boundary and the tangential alignment decreases along the radius, suggesting a complex interaction between the actin network and the inner boundary. For the thicker ring pattern, the actin network almost becomes isotropic (*k*_*H*_~0.65) at the outer boundary. Based on these results, we performed Voronoi cell model simulations of the thick ring-patterned cells (**Figure 6C**). Quantification of the angle deviation of the model cells showed that the outer boundary cells became more radially aligned, larger in area than the innermost cells, and slightly more elongated with increasing condensation tendency and cell stiffness differential (**Figure 6 – figure supplement 2**). We further compared the *in vitro* experimental results with the model prediction (**Figure 7**). We found that for both thick and thin ring patterns, the outermost boundary REF 2c cells radially aligned, but not REF 11b cells or 3T3 cells, which are quantified using angle deviation (**Figure 7**). As predicted in the model, the cell area increased significantly near the pattern boundary for REF 2c (**Figure 7 – figure supplement 1**). Interestingly, the innermost boundary cells align circumferentially along the inner boundary, which is not revealed in our model.

**Figure 6.**
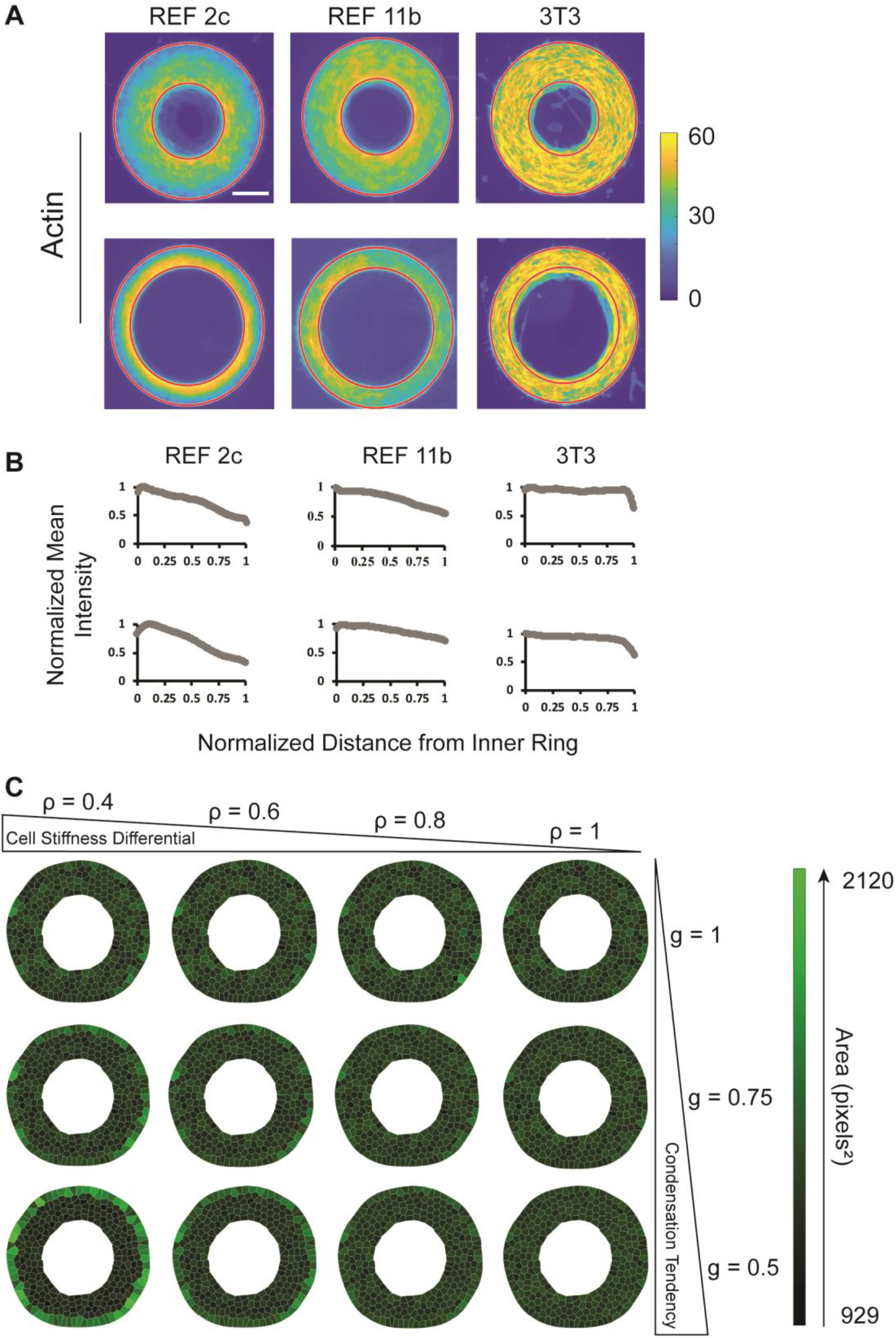
Actin distribution and Voronoi model of cells confined on ring-shaped patterns. (**A**) Heat maps showing actin intensity for REF 2c (*n* = 19 thick ring patterns, *n* = 20 thin ring patterns), REF 11b (*n* = 20 both patterns), and 3T3 (*n* = 20 both patterns). Scale bar: 100 μm. (**B**) Normalized actin intensity is plotted as function of distance from the innermost edge of the rings. (**C**) Voronoi cell ring results predict cell elongation at the outer boundary with large condensation tendency and stiffness differential.

**Figure 7.**
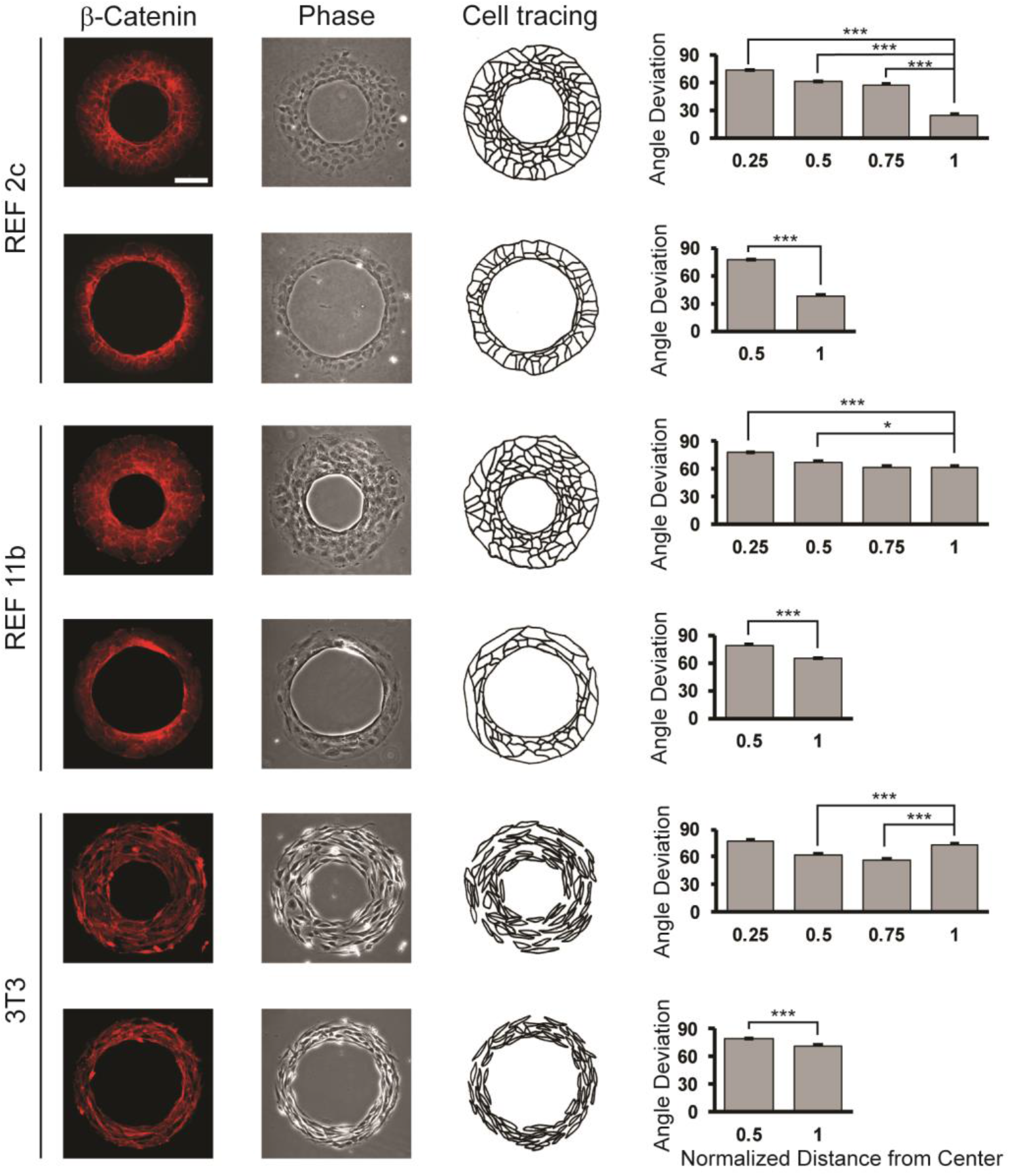
Cell alignment on ring-shaped patterns. Fluorescent images showing β-catenin signals, phase contrast images, and schematics of REF 2c, REF 11b, and 3T3 cells at 48 h. Angle deviation is quantified as a function of normalized distance from center. *n* = 16 patterns per group. Scale bar: 100 μm. Data are represented as mean ± s.e.m. *, *P* < 0.05, ***, *P* < 0.001.

## DISCUSSION

There is a growing interest in understanding physical principles of the autonomous collective behavior of cells. Epithelial cells are usually modelled as a continuum, and the mechanical states are described by the active stress between neighboring cells^4^. On the other hand, actin-rich mesenchymal-like cells are often modelled as self-propelled particles, and they are believed to self-assemble as active nematics^39, 40^. Our results revealed a new class of behavior of the actin-rich cells with significant intercellular adhesion, represented by REF cells. We showed that these cells could form polar symmetry in certain conditions. Our results revealed the REF monolayer displayed a condensation tendency, resulting in a change of cell size and cell elongation. Unlike skeletal muscle cells, such condensation tendency is not originated from the anisotropy of actin bundles, as the actin network in REF monolayer and cell contractility are found to be isotropic (**Figure 4 – figure supplement 1**). Our theoretical analysis suggests that such condensation tendency of the tissue, together with an autonomously established stiffness gradient in the monolayer, are sufficient to form the polar symmetry in the confined planar geometries. Disrupting the cell-cell adhesion using EGTA also abolished the radial cell alignment of REF cells (**Figure 2E**).

In this work, we describe the cell monolayer mechanics associated with the cell contractility and differential cell stiffness by developing a continuum model and a cell-based model. Vertex models^41^ and Voronoi cell models^30, 31^ have been well-established for epithelial-like cells with strong cell-cell adhesion^41^, because of the cortical distribution of actin cytoskeleton. For fiber-rich fibroblast-like cells, current active nematic theory treats individual cells as incompressible, polarized particles. In previous continuum models of monolayer mechanics, the contractility in active nematic theory is described as a traceless stress tensor resulting in anisotropic contraction accompanied by lateral expansion. To better describe the effect of condensation tendency in the REF monolayer, we adopt the morphoelasticity theory where this tendency is described as an isotropic prestretch tensor that induces isotropic area loss in condensation. In previous cell-based modeling, vertex models^41^ and Voronoi cell models^30, 31^ often describe the epithelial cell contractility along the intercellular junctions due to the cortical distribution of actin cytoskeleton with strong cell-cell adhesion. In our case of REF monolayer where the actin network does not spread coincidentally with the intercellular junctions, we modify the Voronoi cell model to describe the condensation tendency as a prestretch on individual cell area, which has not been considered in the previous Voronoi cell models. From both models, we have shown that condensation tendency and a supracellular cell stiffness gradient are sufficient to establish the polarity in the tissue continuum.

The cell system we described here is different from epithelial cells, which also have strong intercellular adhesion and contractility as individual cells. Epithelial cells, when confined in circular patterns, form nematic symmetry that is similar to 3T3 cells^10, 36^. However, when they are confined in ring-shape patterns, chiral spiral defects can be found^42^. The different cell alignment between epithelial cells and REF cells is likely due to the distribution of actin network. It’s well-known that actin mainly distributed in the cell-cell junctions for epithelial cells^43^, while actin stress fibers can be found in REF cells even when they form a compact monolayer. Notably, neuroepithelial cells derived from pluripotent stem cells could form rosettes-like structures *in vitro,* with tight and adherence junctions presented at the side facing the internal lumen^44, 45^. However, the formation of neural rosettes may require a completely different mechanism as neuroepithelial cells are planar polarized^46^, and the junctional proteins distributed homogenously for REF cells. Together, the REF cells represented a unique class of cell systems that are different from typical spindle-like 3T3 cells and epithelial cells. Future studies will be needed to investigate whether other cells that may fall into this category, such as neural crest cells and chondrocytes, also behavior similarly on patterned substrates. Interestingly, neural crest cells migrate as a cohesive group mediated by self-secreted C3a^47^, and chondrocytes form mesenchymal condensation mediated by EGF signals^48^. The condensation tendency of these cells may be relevant to their biological functions by facilitating their orientation towards the signal center.

In summary, this work reports a unique behavior of REF cells that develop polar symmetry when confined in circular and ring mesoscale patterns. The formation of such radial alignment is not a result of directed cell migration and requires a condensation tendency and an emergent supracellular cell stiffness gradient. Future work must be done to understand the molecular mechanisms of this novel collective cell behavior, and its contribution to relevant biological functions.

## ACKNOWLEDGMENT

This work is supported in part by the National Science Foundation (CMMI 1662835 and CMMI 1846866 to Y.S., and DMS 2012330 to M.W), the Department of Mechanical and Industrial Engineering at the University of Massachusetts Amherst, and Department of Mathematical Sciences at Worcester Polytechnic Institute. The Conte Nanotechnology Cleanroom Lab is acknowledged for support in microfabrication. The authors acknowledge the University of Massachusetts Amherst Light Microscopy Core for confocal microscopy and Genomics Resource Laboratory for genomics services.

## AUTHOR CONTRIBUTIONS

T.X., S.S., M.W., and Y.S. designed the experiments, T.X. and S.S. performed *in vitro* experiments, T.X., S.S., N.O., L.B analyzed data. S.S., N.O., M.W. developed the continuum and Voronoi cell models. All authors wrote and approved the manuscript.

## REFERENCES

1. Friedl P, Hegerfeldt Y, Tusch M. Collective cell migration in morphogenesis and cancer. Int J Dev Biol. 2004;48.

2. Gurtner GC, Werner S, Barrandon Y, Longaker MT. Wound repair and regeneration. Nature. 2008;453(7193):314–21.

3. Trepat X, Chen Z, Jacobson K. Cell migration. Comprehensive Physiology. 2012;2(4):2369–92.

4. Trepat X, Sahai E. Mesoscale physical principles of collective cell organization. Nat Phys. 2018;14(7):671–82.

5. Tambe DT, Hardin CC, Angelini TE, Rajendran K, Park CY, Serra-Picamal X, Zhou EHH, Zaman MH, Butler JP, Weitz DA, Fredberg JJ, Trepat X. Collective cell guidance by cooperative intercellular forces. Nat Mater. 2011;10(6):469–75.

6. Trepat X, Fredberg JJ. Plithotaxis and emergent dynamics in collective cellular migration. Trends in Cell Biology. 2011;21(11):638–46.

7. Rodriguez-Franco P, Brugues A, Marin-Llaurado A, Conte V, Solanas G, Batlle E, Fredberg JJ, Roca-Cusachs P, Sunyer R, Trepat X. Long-lived force patterns and deformation waves at repulsive epithelial boundaries. Nat Mater. 2017;16(10):1029−+.

8. Duclos G, Erlenkamper C, Joanny JF, Silberzan P. Topological defects in confined populations of spindle-shaped cells. Nat Phys. 2017;13(1):58–62.

9. Kawaguchi K, Kageyama R, Sano M. Topological defects control collective dynamics in neural progenitor cell cultures. Nature. 2017;545(7654):327−+.

10. Saw TB, Doostmohammadi A, Nier V, Kocgozlu L, Thampi S, Toyama Y, Marcq P, Lim CT, Yeomans JM, Ladoux B. Topological defects in epithelia govern cell death and extrusion. Nature. 2017;544(7649):212–6.

11. Kemkemer R, Kling D, Kaufmann D, Gruler H. Elastic properties of nematoid arrangements formed by amoeboid cells. The European Physical Journal E. 2000;1(2):215–25.

12. Marchetti MC, Joanny JF, Ramaswamy S, Liverpool TB, Prost J, Rao M, Simha RA. Hydrodynamics of soft active matter. Rev Mod Phys. 2013;85(3).

13. Surrey T, Nedelec F, Leibler S, Karsenti E. Physical properties determining self-organization of motors and microtubules. Science. 2001;292(5519):1167–71.

14. Ostrovidov S, Hosseini V, Ahadian S, Fujie T, Parthiban SP, Ramalingam M, Bae H, Kaji H, Khademhosseini A. Skeletal muscle tissue engineering: methods to form skeletal myotubes and their applications. Tissue engineering Part B, Reviews. 2014;20(5):403–36.

15. Achilleos A, Trainor PA. Neural crest stem cells: discovery, properties and potential for therapy. Cell Research. 2012;22(2):288–304.

16. Aomatsu E, Chosa N, Nishihira S, Sugiyama Y, Miura H, Ishisaki A. Cell-cell adhesion through N-cadherin enhances VCAM-1 expression via PDGFR beta in a ligand-independent manner in mesenchymal stem cells. International Journal of Molecular Medicine. 2014;33(3):565–72.

17. Tavella S, Raffo P, Tacchetti C, Cancedda R, Castagnola P. N-CAM and N-Cadherin Expression during in Vitro Chondrogenesis. Experimental Cell Research. 1994;215(2):354–62.

18. Mary S, Charrasse S, Meriane M, Comunale F, Travo P, Blangy A, Gauthier-Rouviere C. Biogenesis of N-cadherin-dependent cell-cell contacts in living fibroblasts is a microtubule-dependent kinesin-driven mechanism. Mol Biol Cell. 2002;13(1):285–301.

19. Takebe T, Enomura M, Yoshizawa E, Kimura M, Koike H, Ueno Y, Matsuzaki T, Yamazaki T, Toyohara T, Osafune K, Nakauchi H, Yoshikawa HY, Taniguchi H. Vascularized and Complex Organ Buds from Diverse Tissues via Mesenchymal Cell-Driven Condensation. Cell Stem Cell. 2015;16(5):556–65.

20. Szabo A, Mayor R. Mechanisms of Neural Crest Migration. Annu Rev Genet. 2018;52:43–63.

21. Piccinini F, Kiss A, Horvath P. CellTracker (not only) for dummies. Bioinformatics. 2016;32(6):955–7.

22. Zhu P, Tseng N-H, Xie T, Li N, Fitts-Sprague I, Peyton SR, Sun Y. Biomechanical Microenvironment Regulates Fusogenicity of Breast Cancer Cells. ACS Biomaterials Science & Engineering. 2019;5(8):3817–27.

23. Xie T, Hawkins J, Sun Y. Traction Force Measurement Using Deformable Microposts. In: Rittié L, editor. Fibrosis: Methods and Protocols. New York, NY: Springer New York; 2017. p. 235–44.

24. Rezakhaniha R, Agianniotis A, Schrauwen JTC, Griffa A, Sage D, Bouten CVC, van de Vosse FN, Unser M, Stergiopulos N. Experimental investigation of collagen waviness and orientation in the arterial adventitia using confocal laser scanning microscopy. Biomech Model Mechanobiol. 2012;11(3-4):461–73.

25. Schneider CA, Rasband WS, Eliceiri KW. NIH Image to ImageJ: 25 years of image analysis. Nat Methods. 2012;9(7):671–5.

26. Ben Amar M, Wu M. Re-epithelialization: advancing epithelium frontier during wound healing. J R Soc Interface. 2014;11(93):20131038.

27. Holzapfel GA. Nonlinear Solid Mechanics: A Continuum Approach for Engineering: Wiley; 2000.

28. Ambrosi D, Ben Amar M, Cyron CJ, DeSimone A, Goriely A, Humphrey JD, Kuhl E. Growth and remodelling of living tissues: perspectives, challenges and opportunities. J R Soc Interface. 2019;16(157).

29. Yan H, Ramirez-Guerrero D, Lowengrub J, Wu M. Stress generation, relaxation and size control in confined tumor growth. bioRxiv. 2019:761817.

30. Bi D, Yang X, Marchetti MC, Manning ML. Motility-driven glass and jamming transitions in biological tissues. Phys Rev X. 2016;6(2).

31. Olaranont N. 2-D Epithelial Tissues, Cell Mechanics, and Voronoi Tessellation: WPI; 2019.

32. Sun Y, Yong KMA, Villa-Diaz LG, Zhang X, Chen W, Philson R, Weng S, Xu H, Krebsbach PH, Fu J. Hippo/YAP-mediated rigidity-dependent motor neuron differentiation of human pluripotent stem cells. Nat Mater. 2014;13(6):599–604.

33. Beningo KA, Hamao K, Dembo M, Wang Y-L, Hosoya H. Traction forces of fibroblasts are regulated by the Rho-dependent kinase but not by the myosin light chain kinase. Arch Biochem Biophys. 2006;456(2):224–31.

34. Ghibaudo M, Saez A, Trichet L, Xayaphoummine A, Browaeys J, Silberzan P, Buguin A, Ladoux B. Traction forces and rigidity sensing regulate cell functions. Soft Matter. 2008;4(9):1836–43.

35. Chen GK, Hou ZG, Gulbranson DR, Thomson JA. Actin-Myosin Contractility Is Responsible for the Reduced Viability of Dissociated Human Embryonic Stem Cells. Cell Stem Cell. 2010;7(2):240–8.

36. Doxzen K, Vedula SRK, Leong MC, Hirata H, Gov NS, Kabla AJ, Ladoux B, Lim CT. Guidance of collective cell migration by substrate geometry. Integrative Biology. 2013;5(8):1026–35.

37. Tavares S, Vieira AF, Taubenberger AV, Araujo M, Martins NP, Bras-Pereira C, Polonia A, Herbig M, Barreto C, Otto O, Cardoso J, Pereira-Leal JB, Guck J, Paredes J, Janody F. Actin stress fiber organization promotes cell stiffening and proliferation of pre-invasive breast cancer cells. Nat Commun. 2017;8:15237.

38. Wu M, Ben Amar M. Modelling fibers in growing disks of soft tissues. Math Mech Solids. 2015;20(6):663–79.

39. Duclos G, Garcia S, Yevick HG, Silberzan P. Perfect nematic order in confined monolayers of spindle-shaped cells. Soft Matter. 2014;10(14):2346–53.

40. Doostmohammadi A, Ignés-Mullol J, Yeomans JM, Sagués F. Active nematics. Nature Communications. 2018;9(1):3246.

41. Farhadifar R, Roper JC, Algouy B, Eaton S, Julicher F. The influence of cell mechanics, cell-cell interactions, and proliferation on epithelial packing. Curr Biol. 2007;17(24):2095–104.

42. Wan LQ, Ronaldson K, Park M, Taylor G, Zhang Y, Gimble JM, Vunjak-Novakovic G. Micropatterned mammalian cells exhibit phenotype-specific left-right asymmetry. Proc Natl Acad Sci U S A. 2011;108(30):12295–300.

43. Fanning AS, Van Itallie CM, Anderson JM. Zonula occludens-1 and-2 regulate apical cell structure and the zonula adherens cytoskeleton in polarized epithelia. Mol Biol Cell. 2012;23(4):577–90.

44. Knight GT, Lundin BF, Iyer N, Ashton LMT, Sethares WA, Willett RM, Ashton RS. Engineering induction of singular neural rosette emergence within hPSC-derived tissues. Elife. 2018;7.

45. Elkabetz Y, Panagiotakos G, Al Shamy G, Socci ND, Tabar V, Studer L. Human ES cell-derived neural rosettes reveal a functionally distinct early neural stem cell stage. Gene Dev. 2008;22(2):152–65.

46. Davey CF, Mathewson AW, Moens CB. PCP Signaling between Migrating Neurons and their Planar-Polarized Neuroepithelial Environment Controls Filopodial Dynamics and Directional Migration. Plos Genet. 2016;12(3).

47. Carmona-Fontaine C, Theveneau E, Tzekou A, Tada M, Woods M, Page KM, Parsons M, Lambris JD, Mayor R. Complement Fragment C3a Controls Mutual Cell Attraction during Collective Cell Migration. Developmental Cell. 2011;21(6):1026–37.

48. Qin L, Beier F. EGFR Signaling: Friend or Foe for Cartilage? JBMR Plus. 2019;3(2):e10177.

