## Supplementary material for "Condensation tendency of connected contractile tissue with planar isotropic actin network": Figure Supplements

**Figure 1, figure supplement 1**

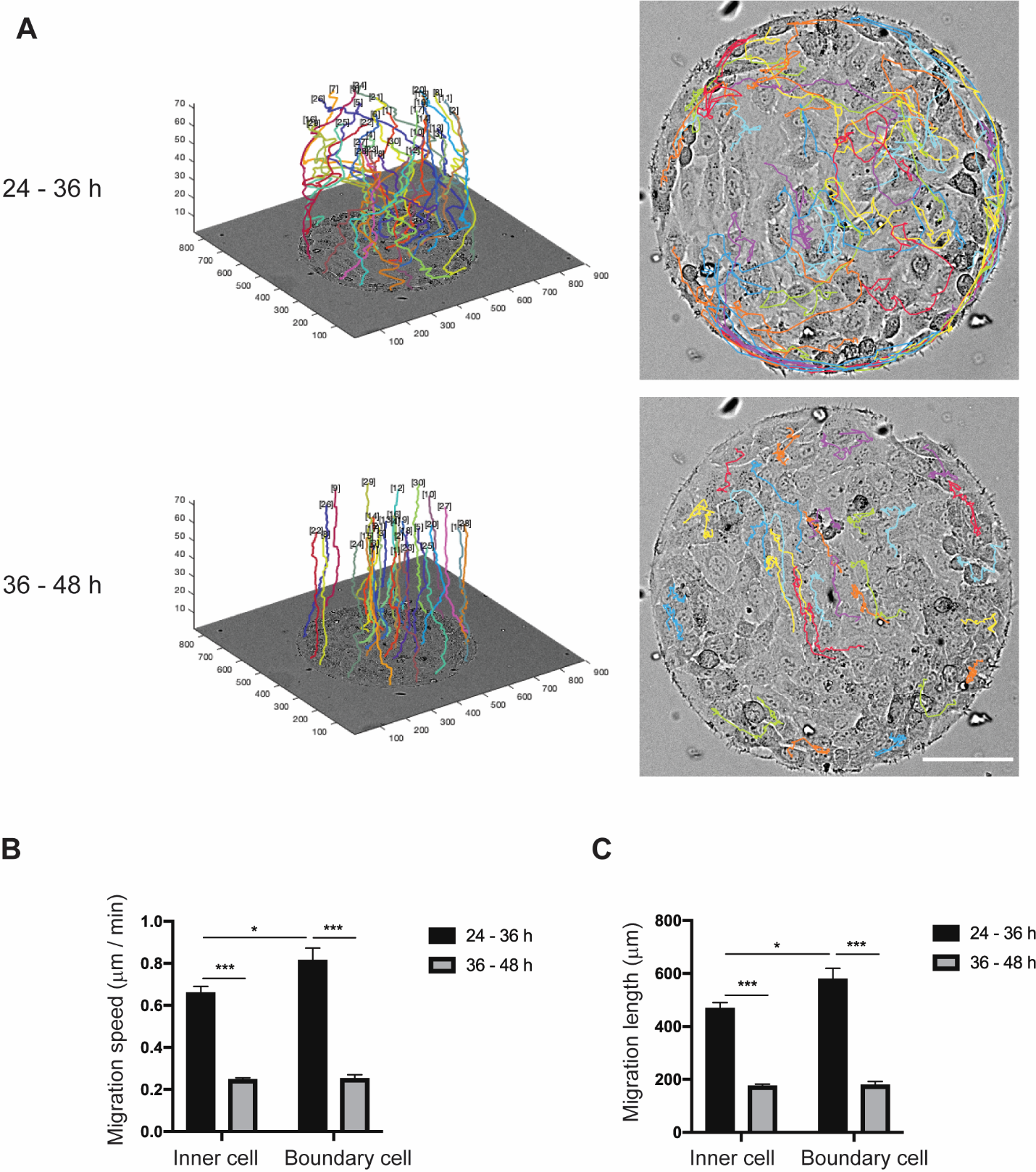

**Figure 1 - figure supplement 1.** Migration of REF 2c cells on circular patterns. (**A**) The 3D and 2D migration trajectories of REF 2c cells (*n* = 30) over 24 - 36 h and 36 - 48 h after seeding. Scale bar, 100 μm. (**B-C**) Migration speed (**B**) and migration length (**C**) of inner (*n* = 15) and boundary cells (*n* = 15) over 24 - 36 h and 36 - 48 h after seeding. Data are presented as mean ± s.e.m. *, *P* < 0.05; ***, *P* < 0.001.

**Figure 1, figure supplement 2**

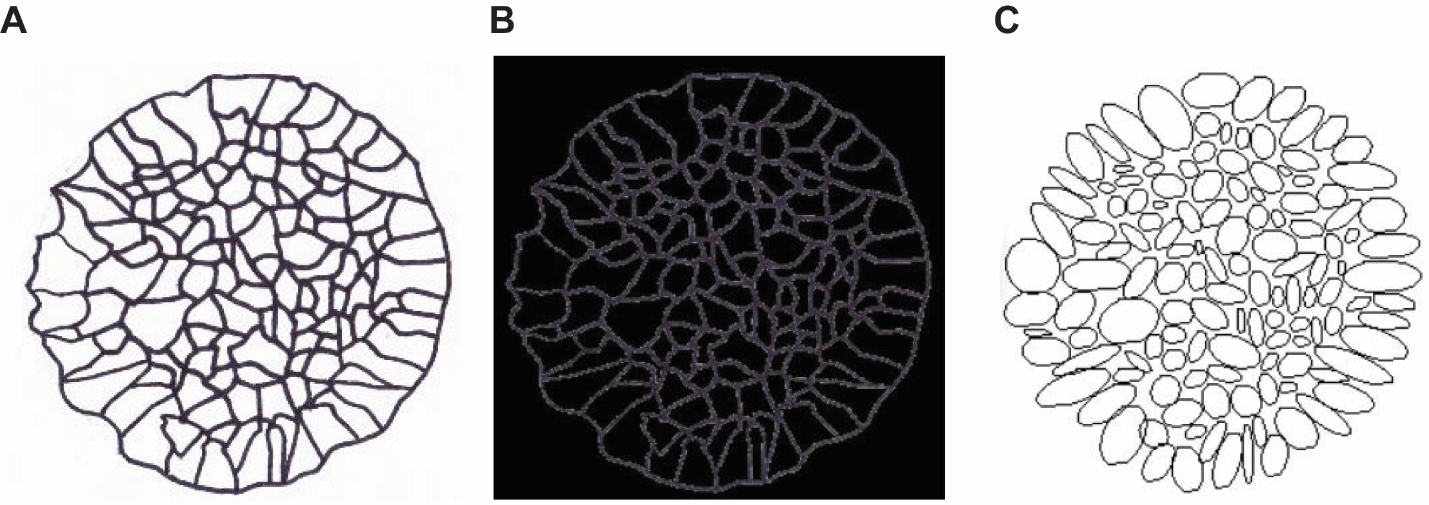

**Figure 1 - figure supplement 2.** Image analysis method for the quantification of angle deviation, cell elongation, and cell area. (**A**) Each cell from the phase contract images was hand-traced to produce a clear outline. (**B**) Tracing was imported into ImageJ and thresholded to create a black background with white cell outlines. (**C**) Ellipses were created using the “analyze particles” tool. Location of pattern centroid was also measured using the circle fit tool. Ellipse attributes (centroid, angle, axes lengths, area) combined with the pattern centroid were used to find cell angle deviation from the radial direction, cell area, and cell elongation as a function of distance from the center of the pattern.

**Figure 1, figure supplement 3**

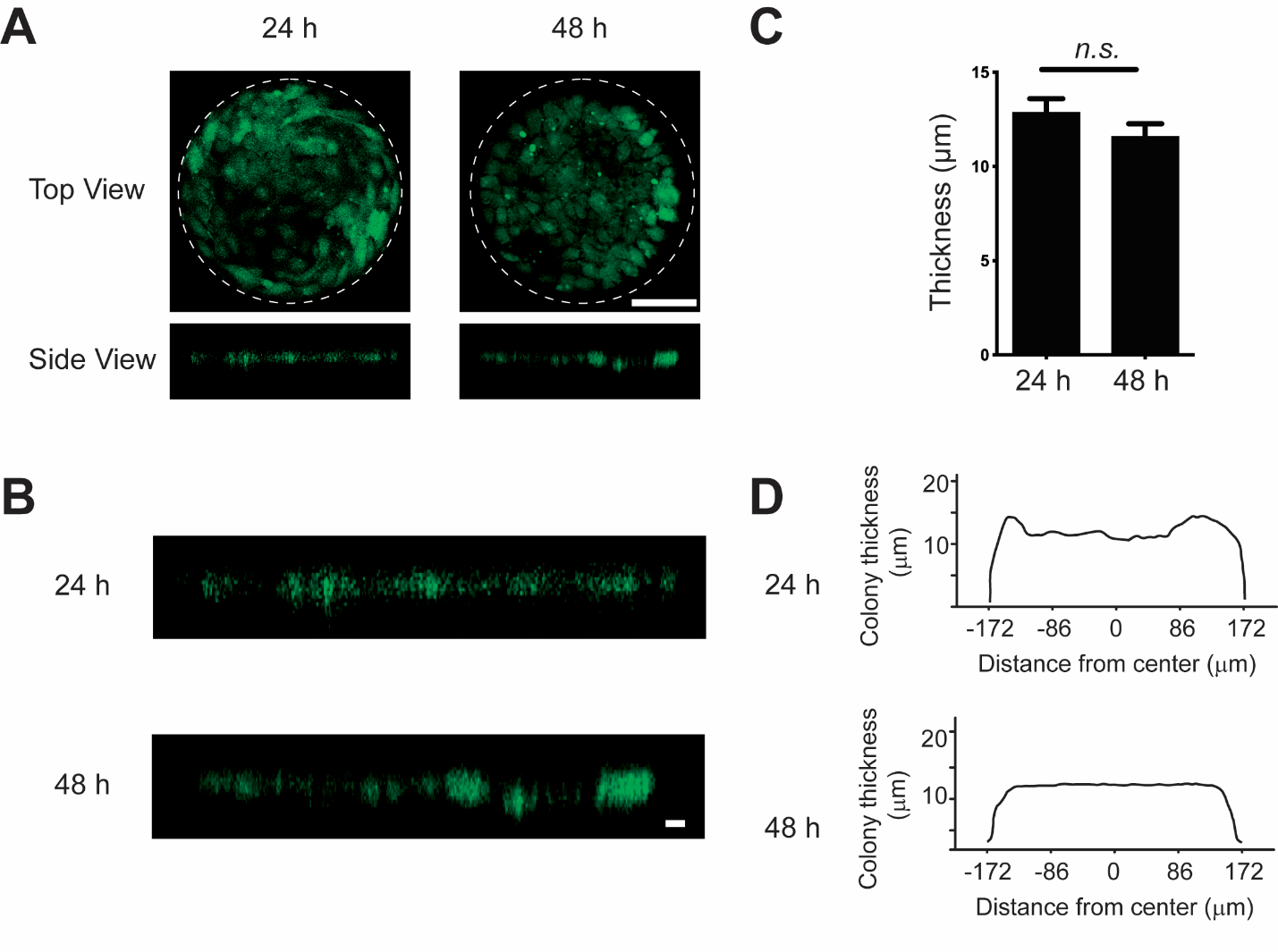

**Figure 1 - figure supplement 3.** The thickness of micropatterned REF 2c colony does not change significantly from 24 h to 48 h. (**A**) The top view and side view of REF 2c labelled with CellTracker-Green dye at 24 h and 48 h. Scale bar, 100 μm. (**B**) Zoomed side view of REF 2c labelled with CellTracker-Green dye at 24 h and 48 h. Scale bar, 10 μm. (C) Bar plot showing the tissue thickness at 24 h and 48 h. *n.s.*, *P* > 0.05. (**D**) Plot showing the mean thickness across the REF colony at 24 h (*n* = 6) and 48 h (*n* = 6).

**Figure 1, figure supplement 4**

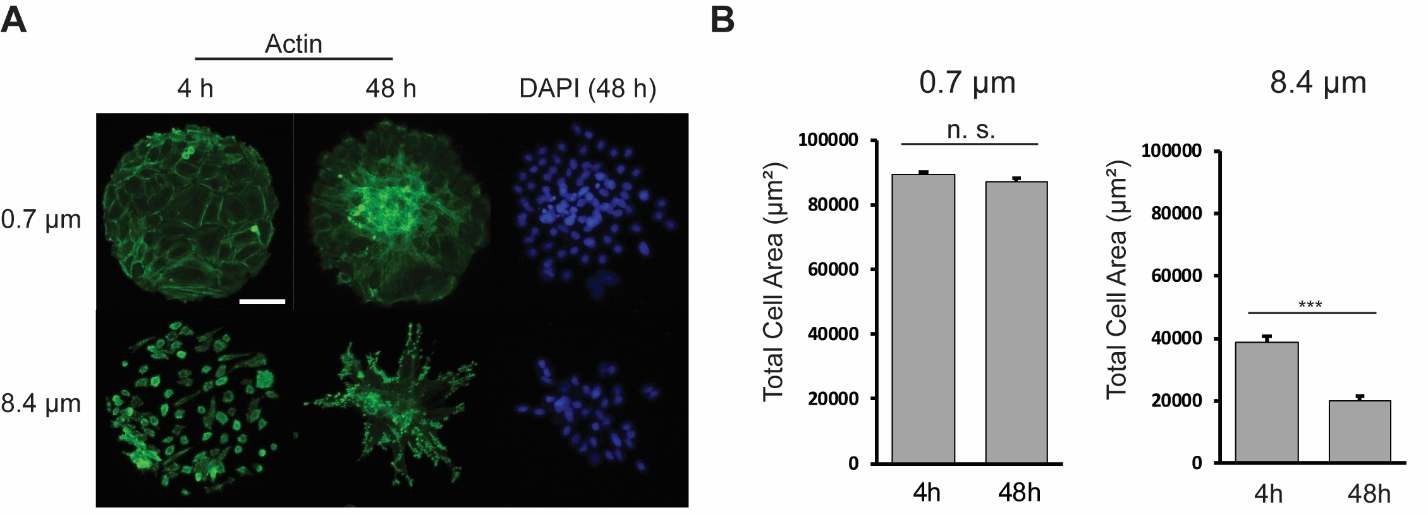

**Figure 1 – figure supplement 4.** Effect of substrate stiffness on the cell alignment of REF 2c cells. (**A**) Representative actin (green) and DAPI (blue) images of REF 2c on PMA substrates with different post heights. Cells were fixed and imaged at 4 h and 48 h, respectively. Scale bar: 100 μm. (**B**) Total cell area on each micropattern as functions of time and substrate stiffness. *n* = 20 patterns for each condition. Data are presented as mean ± s.e.m. ***, *P* < 0.001; *n.s.*, *P* > 0.05.

**Figure 2, figure supplement 1**

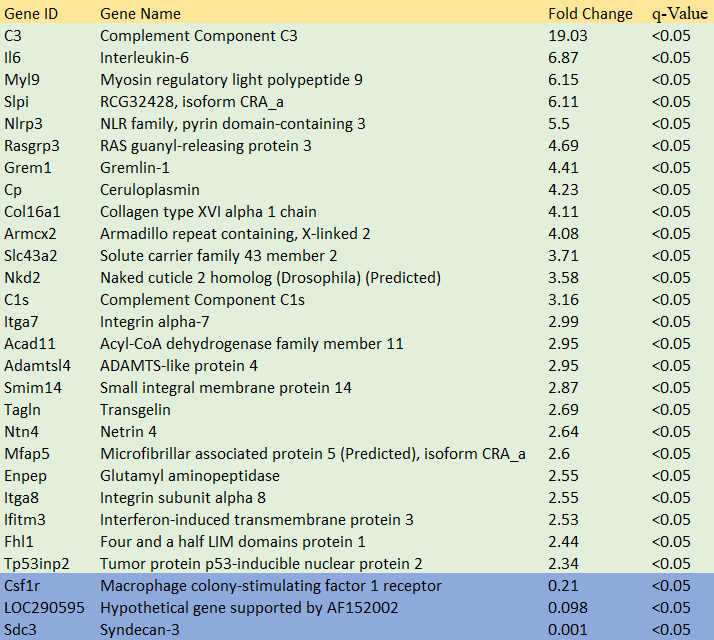

**Figure 2 – figure supplement 1**. Summary of gene expression differences in REF 11b compared to REF 2c.

**Figure 4, figure supplement 1**

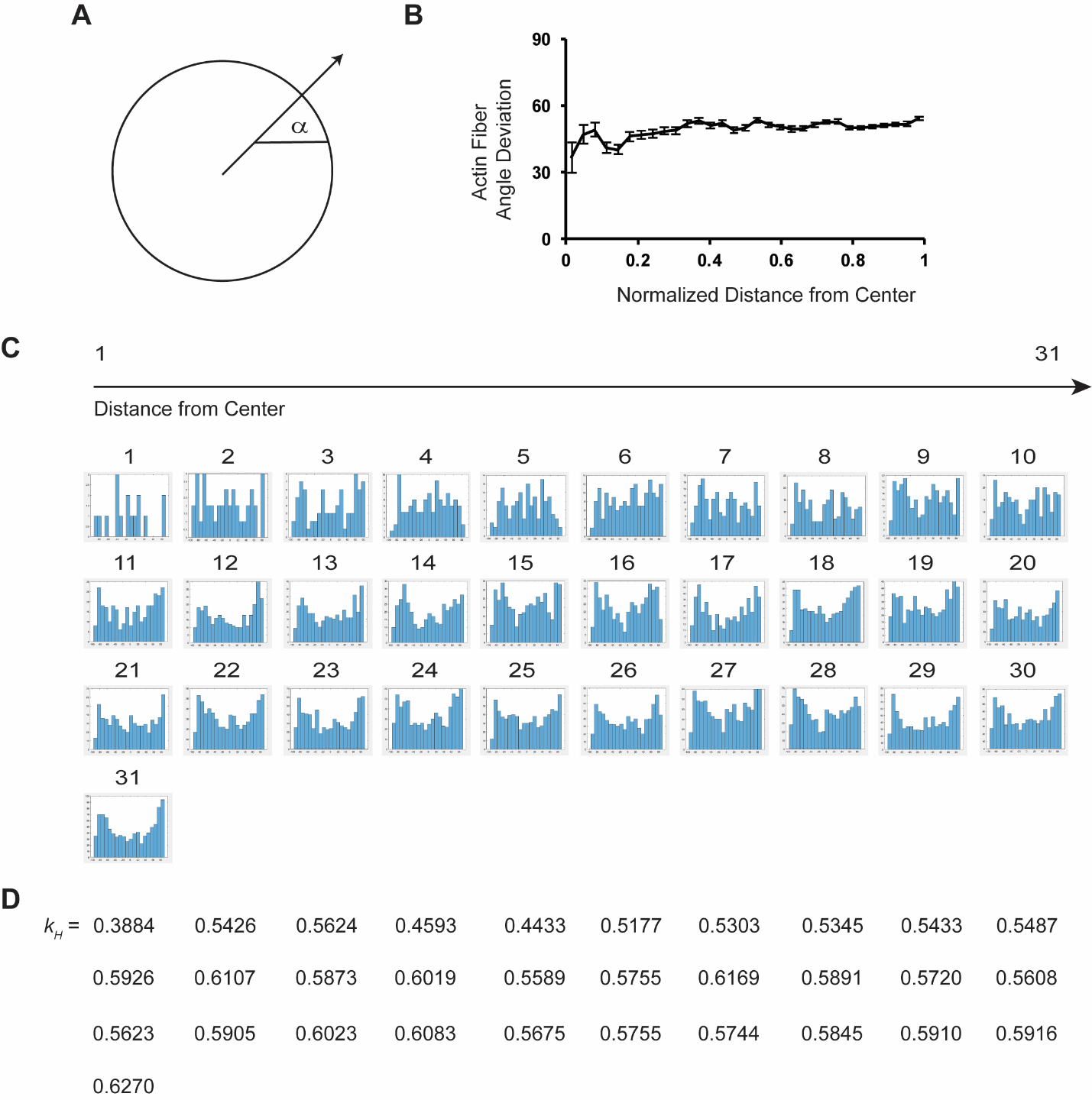

**Figure 4 - figure supplement 1**. Structure tensor developed from histograms of fiber angle deviation. (**A**) Schematic of fiber angle deviation calculation. (**B**) Actin fiber angle deviation as a function of normalized distance from center of the pattern. Data is presented as mean ± s.e.m. (**C**) Histograms of the actin fiber orientation distribution as a function of distance from the center of the pattern. *n* = 5 patterns. (**D**) Structure parameter (*k_H_*) derived from the histograms. *n* = 5 patterns.

**Figure 5, figure supplement 1**

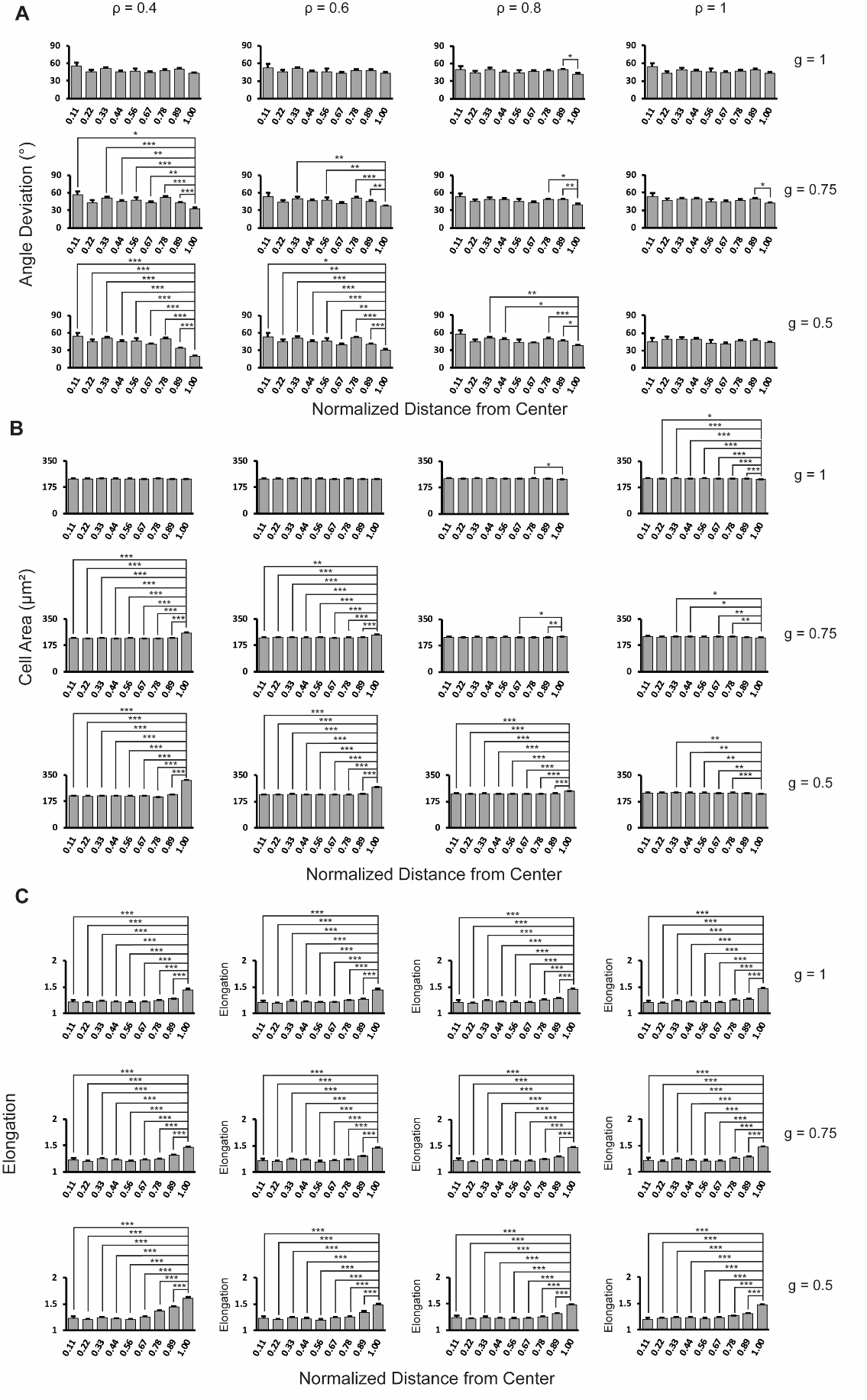

**Figure 5 - figure supplement 1**. Quantitative analysis of the Voronoi cell modeling for circularly patterned cells. (**A**) Angle deviation is plotted as a function of distance from the center of the pattern. *n* = 5 patterns. (**B**) Cell area is plotted as a function of distance from the center of the pattern. *n* = 5 patterns. (**C**) Cell elongation is plotted as a function of distance form the center of the pattern. *n* = 5 patterns. Data are represented as mean ± s.e.m. *, *P* < 0.05, **, *P* < 0.01, ***, *P* < 0.001.

**Figure 6, figure supplement 1**

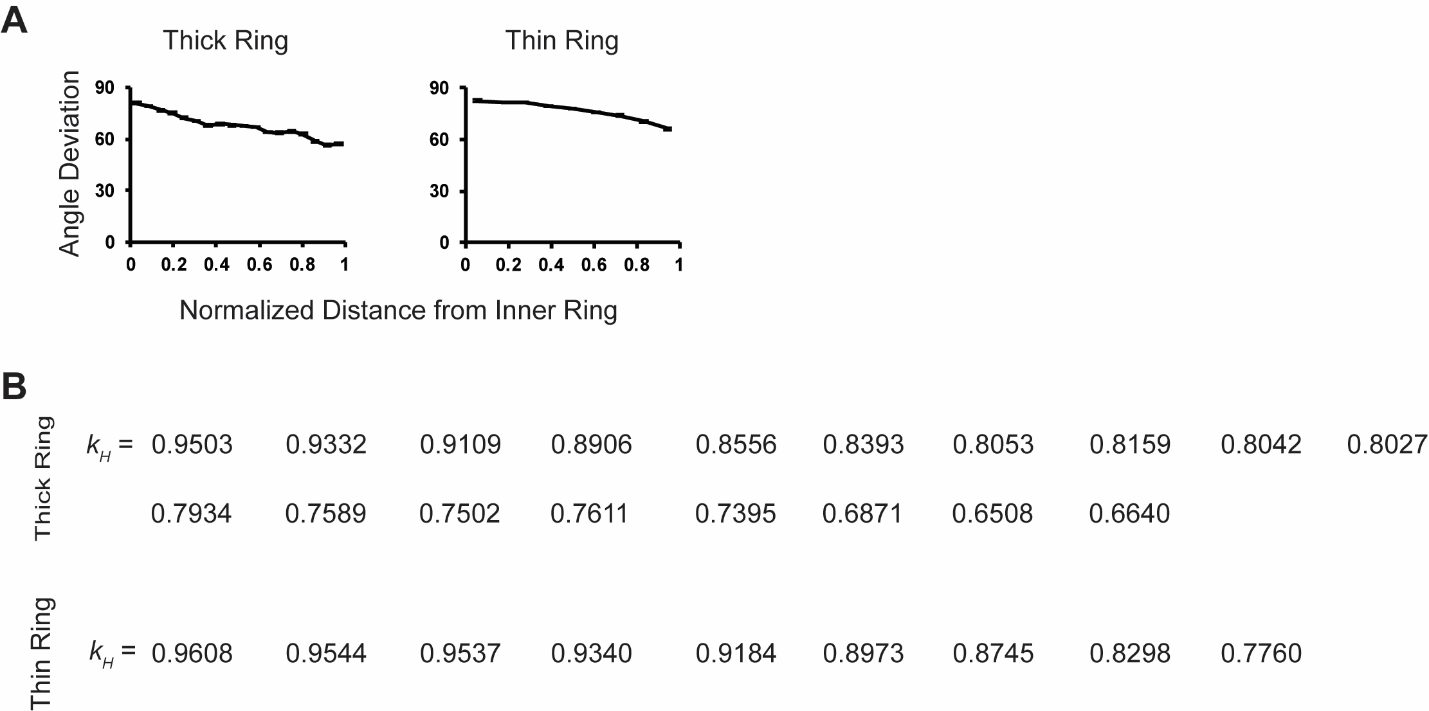

**Figure 6 - figure supplement 1**. Modeling REF 2c cells patterned on ring-shape micropatterns. **(A)** Actin fiber angle deviation is plotted as a function of normalized distance from the center of the innermost edge of the rings. *n* = 5 patterns. (**B**) Structure parameter *k_H_* at derived from the histograms of actin fiber orientation for thick and thin rings. *n* = 5 patterns.

**Figure 6, figure supplement 2**

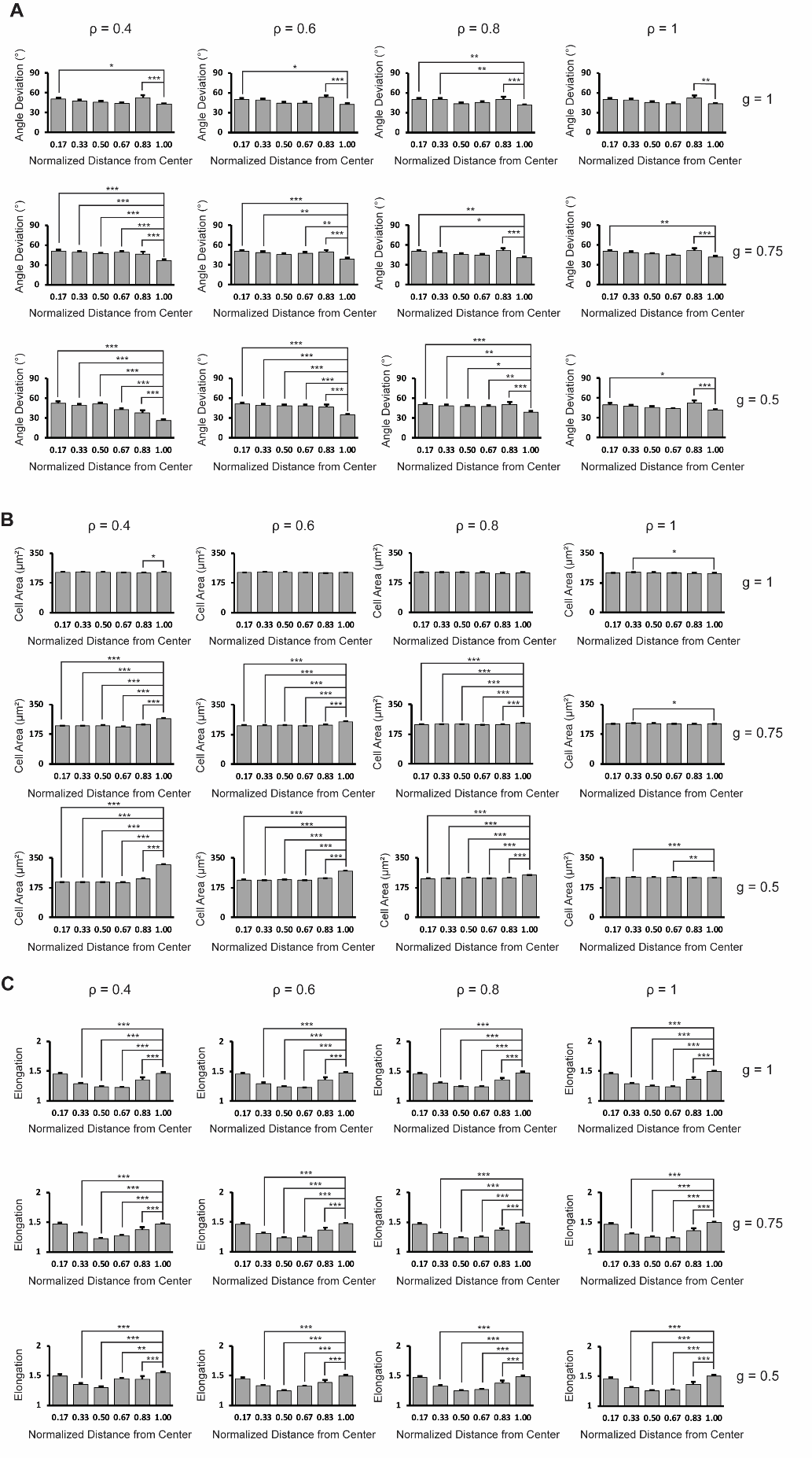

**Figure 6 - figure supplement 2**. Quantitative analysis of the Voronoi cell modeling for ring-patterned cells. (**A**) Angle deviation is plotted as a function of distance from the center of the pattern. *n* = 5 patterns. (**B**) Cell area is plotted as a function of distance from the center of the pattern. *n* = 5 patterns. (**C**) Cell elongation is plotted as a function of distance from the center of the pattern. *n* = 5 patterns. Data are represented as mean ± s.e.m. *, *P* < 0.05, **, *P* < 0.01, ***, *P* < 0.001.

**Figure 7, figure supplement 1**

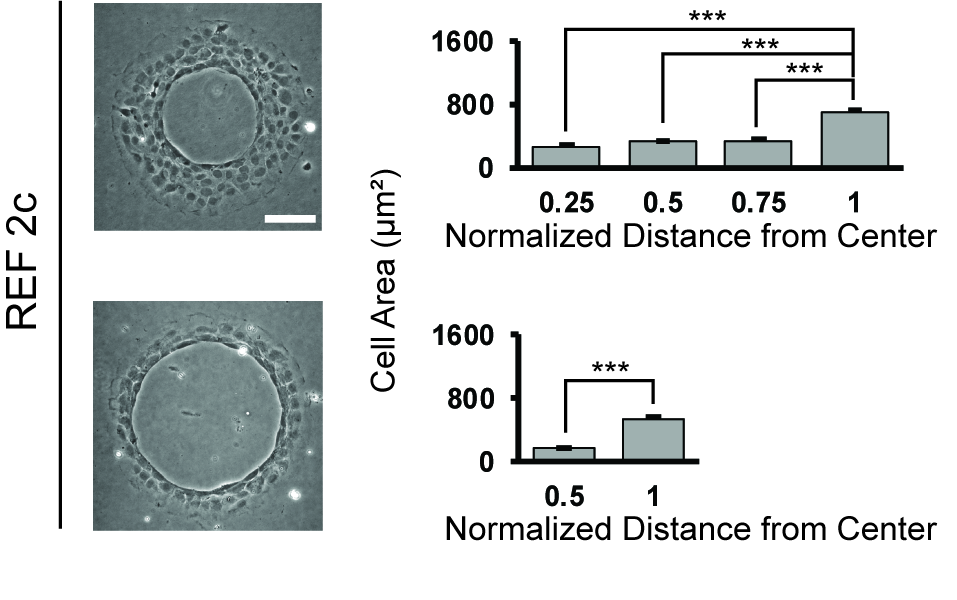

**Figure 7 - figure supplement 1**. Cell area of REF2c cells on ring patterns. Cell area is plotted as a function of distance from the center of the pattern for thick and thin rings. Data are represented as mean ± s.e.m. ***, *P* < 0.001.
